# Viral spillover risk increases with climate change in High Arctic lake sediments

**DOI:** 10.1101/2021.08.23.457348

**Authors:** Audrée Lemieux, Graham A. Colby, Alexandre J. Poulain, Stéphane Aris-Brosou

## Abstract

The host spectrum of viruses is quite diverse, as they can sustainedly infect a few species to several phyla. When confronted with a new host, a virus may even infect it and transmit sustainably in this new host, a process called “viral spillover.” However, the risk of such events is difficult to quantify. As climate change is rapidly transforming environments, it is becoming critical to quantify the potential for spillovers. To address this issue, we resorted to a metagenomics approach and focused on two environments, soil and lake sediments from Lake Hazen, the largest High Arctic freshwater lake in the world. We used DNA and RNA sequencing to reconstruct the lake’s virosphere in both its sediments and soils, as well as its range of eukaryotic hosts. We then estimated the spillover risk by measuring the congruence between the viral and the eukaryotic host phylogenetic trees, and show that spillover risk is higher in lake sediments than in soil, and increases with runoff from glacier melt, a proxy for climate change. Should climate change also shift species range of potential viral vectors and reservoirs northwards, the High Arctic could become fertile ground for emerging pandemics.

---

Viruses are ubiquitous and are often described as the most abundant replicating entities on Earth (Dávila-Ramos et al., 2019; Koonin et al., 2015; Y. Z. Zhang et al., 2018). In spite of having highly diverse genomes, viruses are not independent “organisms” or replicators (Dawkins, 1989), as they need to infect a host’s cell in order to replicate. These virus/host relationships seem relatively stable within superkingdoms, and can hence be classified as *archaeal, bacterial* (also known as *bacteriophages*), and *eukaryotic* viruses (Malik et al., 2017; Nasir et al., 2014, 2017). However, below this rank, viruses may infect a novel host from a reservoir host by being able to transmit sustainably in this new host, a process that, following others (Alexander et al., 2018; Becker et al., 2019), we define as viral spillover. Indeed, in the past years, many viruses such as the Influenza A (Long et al., 2019), Ebola (Rewar & Mirdha, 2014), and SARS-CoV-2 (Zhou & Shi, 2021) spilled over to humans and caused significant diseases. While these three viruses have non-human wild animal reservoirs as natural hosts, others have a broader host range, or their reservoir is more challenging to identify. For instance, iridoviruses are known to infect both invertebrates and vertebrates (İnce et al., 2018), and *Picornavirales* are found in vertebrates, insects, plants, and protists (Koonin et al., 2015). Such a versatile host specificity is difficult to assess without resorting to an expert opinion (Grange et al., 2021), and gauging the probability that a virus may spillover from one host species to another, *i*.*e*., its spillover risk, is hard to quantify.

Numerous factors can influence such a viral spillover risk. For instance, viral particles need to attach themselves to specific receptors on their host’s cell to invade it (Longdon et al., 2014; Maginnis, 2018; Parrish et al., 2008). The conservation of those receptors across multiple species allows these hosts to be more predisposed to becoming infected by the same virus (Parrish et al., 2008; Woolhouse et al., 2005). Indeed, from an evolutionary standpoint, viruses are more prone to infecting hosts that are phylogenetically close to their natural host (Longdon et al., 2014, 2018), potentially because it is easier for them to infect and colonise species that are genetically similar (Geoghegan et al., 2017). Alternatively, but not exclusively, high mutation rates might explain why RNA viruses spill over more often than other viruses (Longdon et al., 2014), as most lack proofreading mechanisms, making them more variable and likely to adapt to a new host (Parrish et al., 2008). RNA viruses are not only more likely to change host, but may do so in novel host species that have different ecological niches as well (Wells et al., 2020).

In this context of spillover, the High Arctic is of special interest as it is particularly affected by climate change, warming faster than the rest of the world (Colby et al., 2020; Lehnherr et al., 2018; Ruuskanen et al., 2020; Meredith et al., 2019). Indeed, a warming climate and rapid transitions of the environment may both increase spillover risk by varying the global distribution and dynamics of viruses, as well as that of their reservoirs and vectors (Parkinson et al., 2014; Waits et al., 2018), as shown for arboviruses (Ciota & Keyel, 2019) and the Hendra virus (Martin et al., 2018). Furthermore, as the climate changes, the metabolic activity of the Arctic’s microbiosphere also shifts, which in turns affects numerous ecosystem processes such as the emergence of new pathogens (Messan et al., 2020). Thus, it has now become critical to be more proactive in preventing such events (**?**), but also to be able to quantify the risk of these spillovers. An intuitive approach to do this is to focus on the cophylogenetic relationships between viruses and their hosts (Bellec et al., 2014; Bennett et al., 2020; Jackson & Charleston, 2004; Madinda et al., 2016; Olival et al., 2017). Conceptually, if both viruses and their hosts cospeciate, the topologies of their respective phylogenetic trees should be identical or *congruent*. On the other hand, the occurrence of spillovers would result in incongruent virus/host phylogenies, so it can be postulated that combining current knowledge about host range and phylogenetic incongruency can be used to quantify spillover risk.

To test this hypothesis in the context of a changing High Arctic environment, we resorted to a combination of metagenomics and of cophylogenetic modelling by sampling both the virosphere and its range of hosts (Y. Z. Zhang et al., 2018), focusing on eukaryotes, which are critically affected by viral spillovers (Carrasco-Hernandez et al., 2017). We contrasted two local environments, lake sediments and soil samples of Lake Hazen, to test how viral spillover risk is affected by glacier runoff, and hence potentially by global warming, which is expected to increase runoff with increasing glacier melt at this specific lake (Colby et al., 2020; Lehnherr et al., 2018). We show here that spillover risk increases with warming climate in lake sediments only, and suggest potential mechanisms explaining these differences.

## Results and Discussion

### Plant and fungal viruses are overrepresented

Based on our most sensitive annotation pipeline (see Electronic Supplementary Material), viruses represented less than 1% of all contigs, and our samples were dominated by bacteria, with low proportions of eukaryotes (proportions of bacterial and eukaryotic contigs being respectively ≥ 89.2% and ≤ 6.4%, in 11 out of 12 samples; see Electronic Supplementary Material). These results could be due to our extraction process, which might have been biased towards microbial nucleic acids, or simply to the fact that bacteria are the major members of soil and sediment samples (Suttner et al., 2020).

Bacteriophages, eukaryotic viruses, and even one virophage species were found among the viral HSPs (see Electronic Supplementary Material). Focusing on eukaryotic viruses, those were found to be unevenly distributed between RNA and DNA genomes, with the former (*i*.*e*., dsRNA, +ssRNA, and -ssRNA viruses) representing between 73.9% to 100% of all hits against eukaryotic viruses (table 1, figure 2). This is not unexpected, as (i) fungal biomass for instance surpasses that of bacteria in Arctic environments by 1-2 orders of magnitude (Schmidt & Bölter, 2002), and (ii) eukaryotes are known to be the main targets of RNA viruses (Koonin et al., 2015; Malik et al., 2017; Nasir et al., 2014, 2017).

**Table 1.**
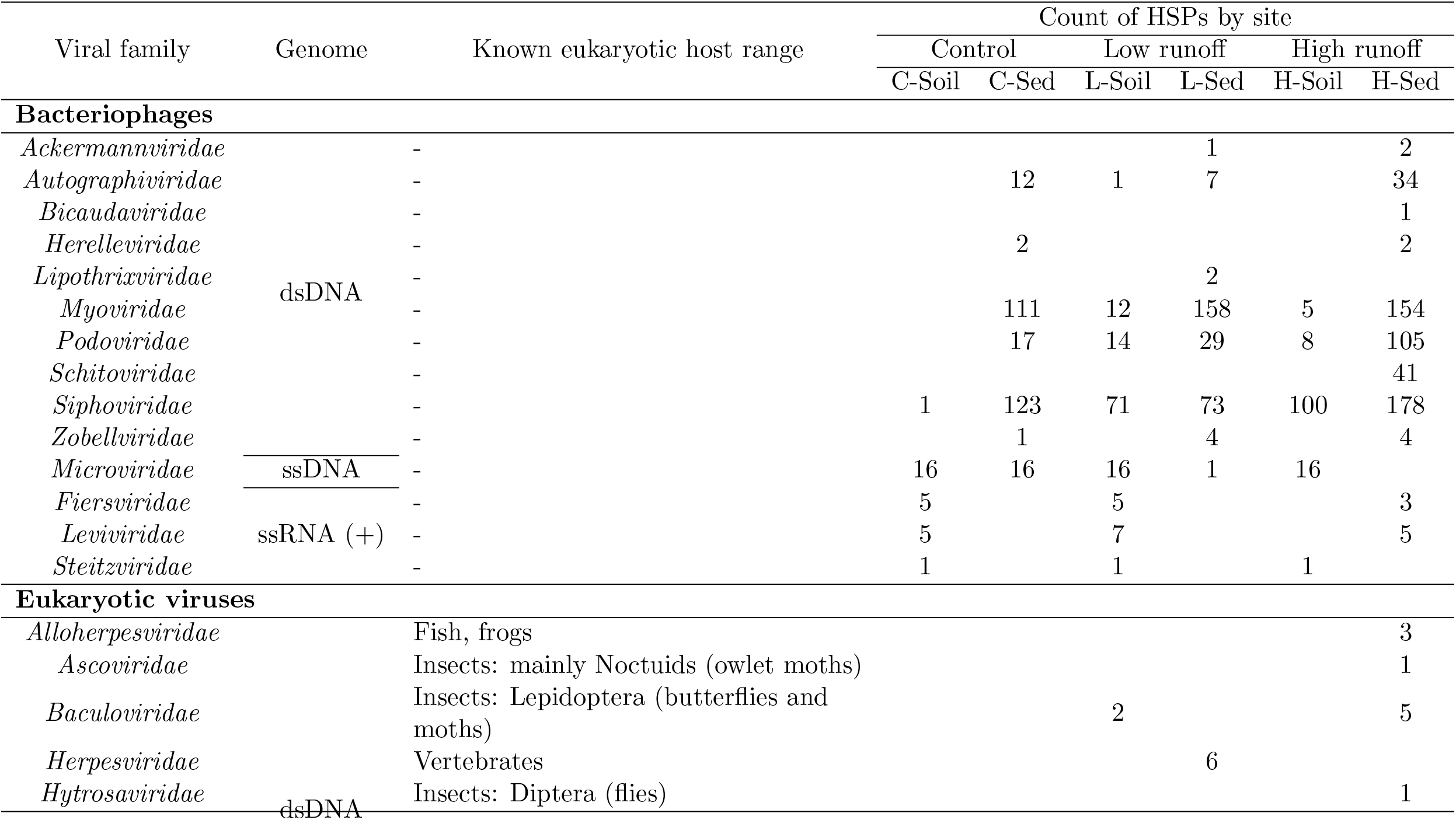
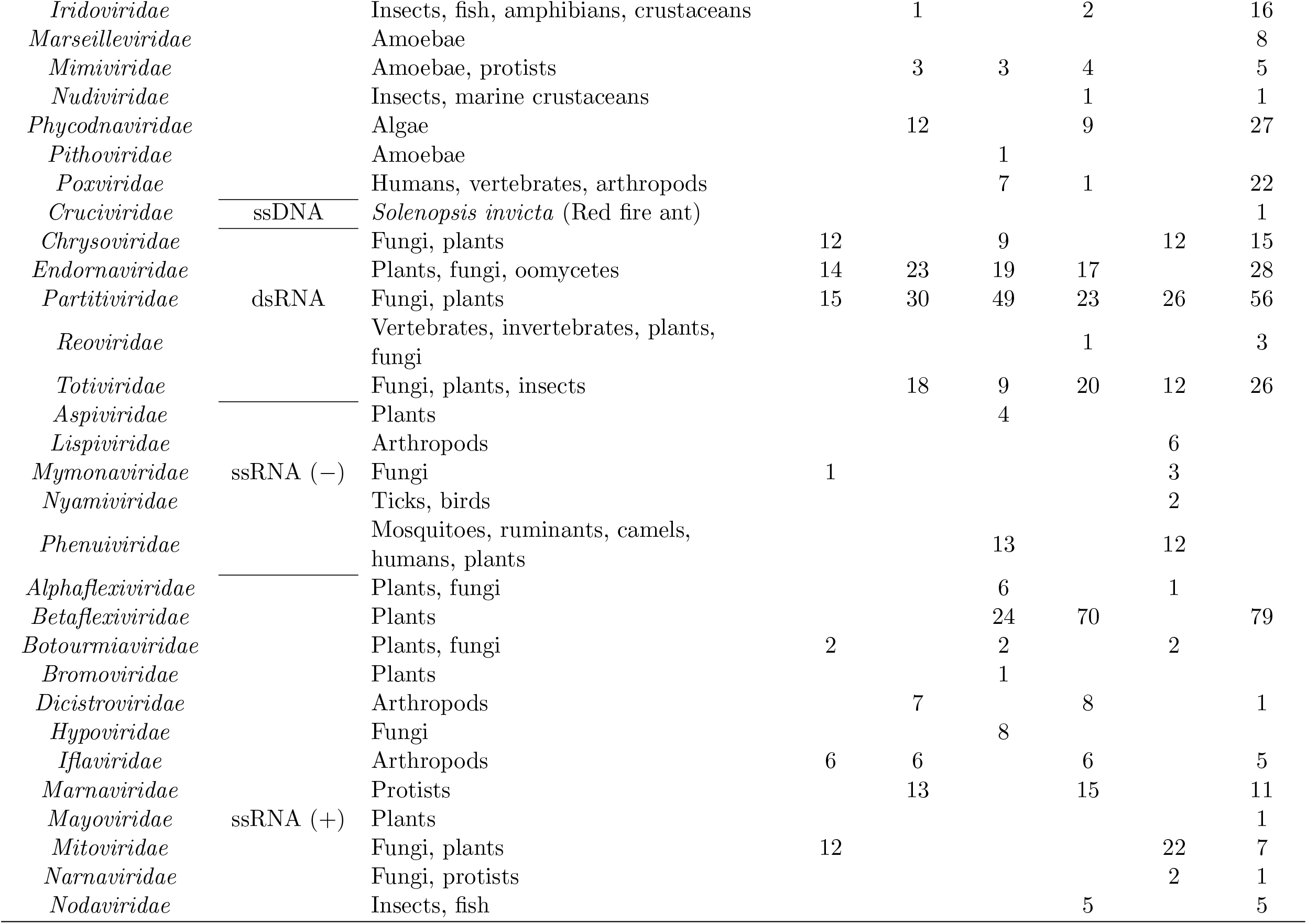
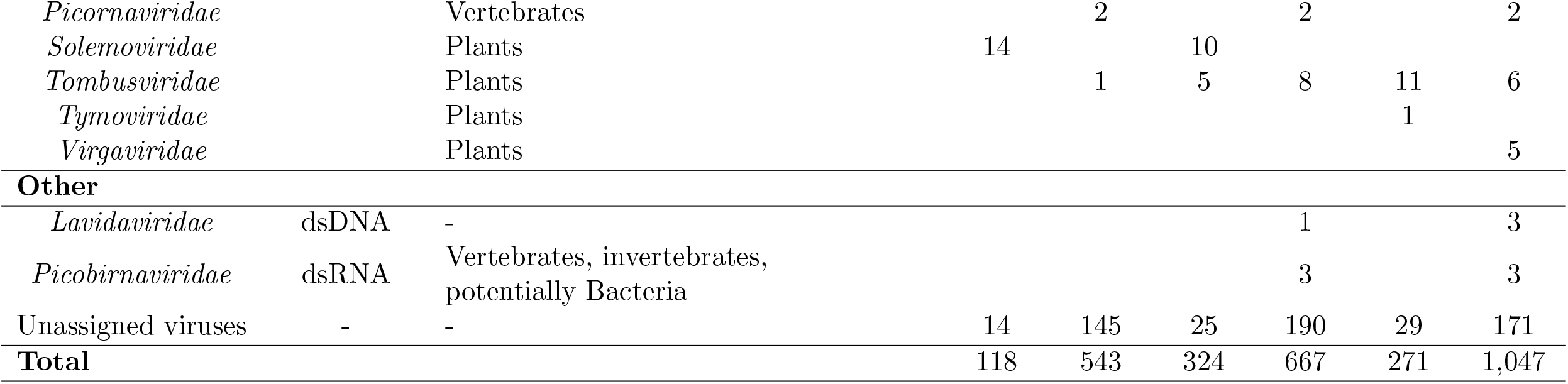
Distribution of the viral families of the viral High-scoring Segment Pairs (HSPs). The host range information was obtained from the ViralZone (Hulo et al., 2011), the International Committee on Taxonomy of Viruses (ICTV) (Lefkowitz et al., 2018), and the Virus-Host (Mihara et al., 2016) databases. *Lavidaviridae* and *Picobirnaviridae* were neither bacteriophages nor eukaryotic viruses, as the former was shown to be virophages (Fischer, 2021) and the latter was recently proposed to be both bacteriophages and eukaryotic viruses (Ghosh & Malik, 2021). Viruses with no or unknown information at the family rank were classified as “Unassigned viruses.” Each viral species was kept once.

**Figure 1.**
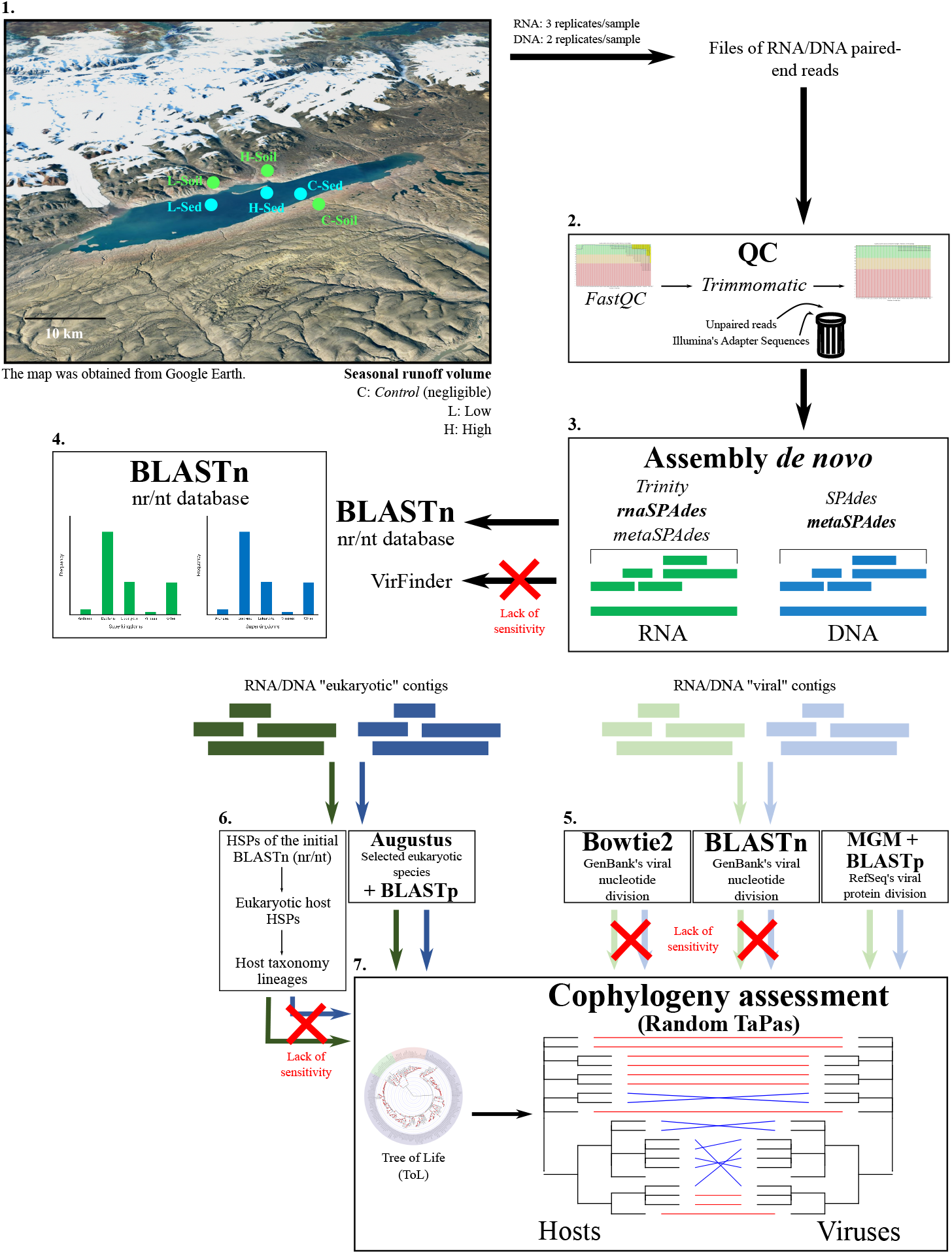
Overview of our analytical pipeline. The following steps are represented: 1. Sampling; 2. Quality control (QC); 3. Assembly *de novo*; 4. Contig classification; 5. Refining of the viral taxonomic annotations; 6. Determination of the eukaryotic hosts from the eukaryotic contigs; and 7. Cophylogeny assessment. DNA and RNA are colour-coded in blue and green, respectively.

**Figure 2.**
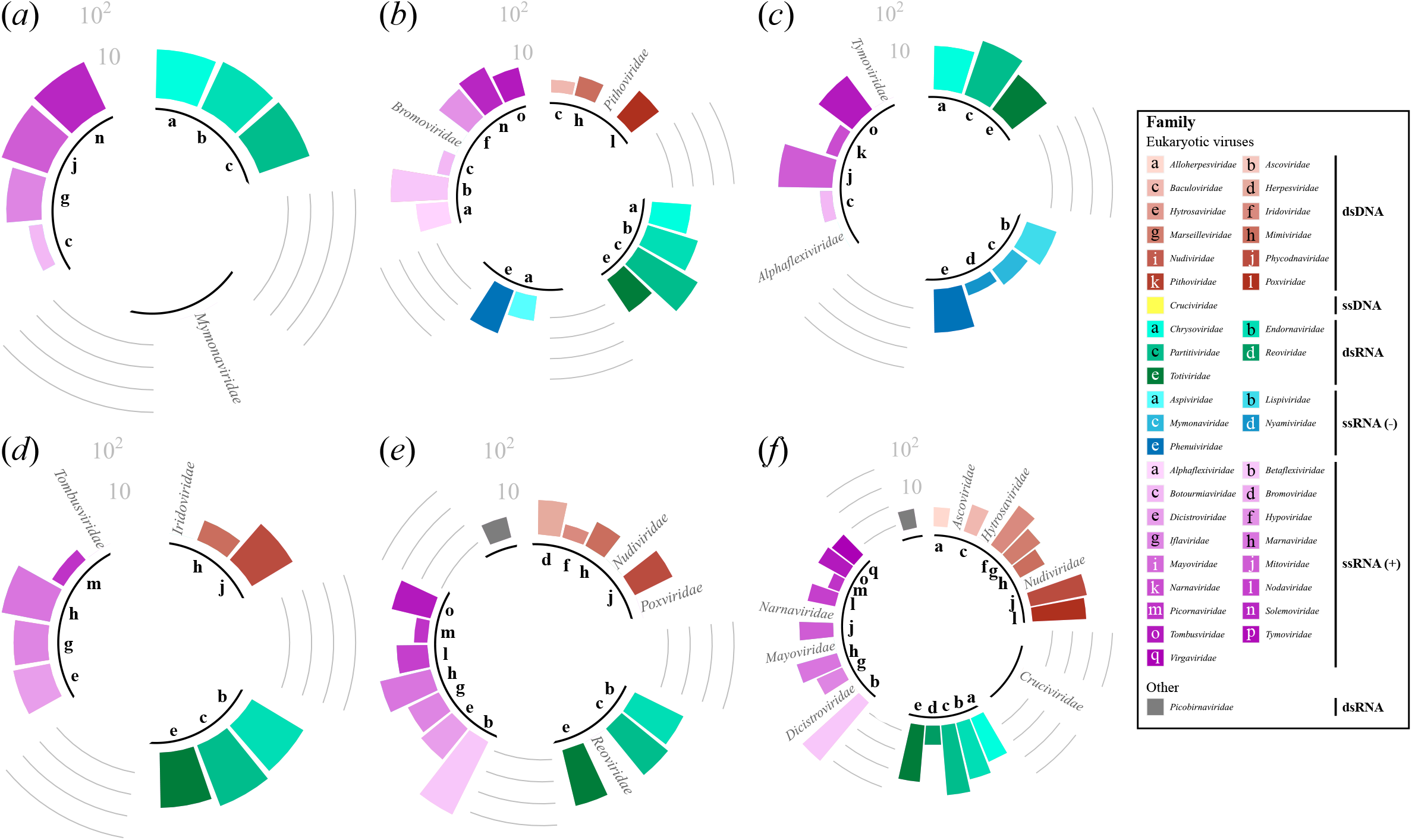
Distribution of the High-scoring Segment Pairs (HSPs) of the eukaryotic viruses, at the family rank. (*a*) C-Soil; (*b*) L-Soil; (*c*) H-Soil; (*d*) C-Sed; (*e*) L-Sed; and (*f*) H-Sed sites. Each viral species was kept once. Viruses of the family *Picobirnaviridae* were neither bacteriophages nor eukaryotic viruses, as they were recently proposed to be both bacteriophages and eukaryotic viruses (Ghosh & Malik, 2021). Counts were log_10_-transformed. The data used for this figure can be found in table 1, and figure S7 depicts the distribution of all viral families, including bacteriophages.

All genomes and samples confounded, the majority of eukaryotic viruses was mainly found to be targeting plants and fungi, with proportions ranging from 62.1% to 92.4% (binomial tests, *P <* 5.93 × 10^*−*3^; table 1). This overrepresentation might reflect a preservation bias, due to the constitutive defences found in plants and fungi offered by their waxy epidermal cuticles and cell walls (Kozieł et al., 2021), even if most plant viruses lack a protective lipoprotein envelope as found in animal viruses (Stavolone & Lionetti, 2017). It is also possible that, in an under-explored environment, virus/host associations not seen in better characterised environments exist, leading us to wrongly associate a virus to a given host, just because unknown associations cannot be learned from what is known. However, there is no reason to believe that such a knowledge gap would tip the bias towards plants and fungi hosts. Irrespective of such a preservation bias, this imbalance could imply a high spillover potential among plants and fungi in the High Arctic for two reasons. First, RNA viruses are the most likely pathogens to switch hosts, due to their high rates of evolution (Longdon et al., 2014; Aris-Brosou et al., 2019). Second, plant biomass has been increasing over the past two decades in the High Arctic due to regional warming (Hudson & Henry, 2009), and is likely to keep doing so as warming continues.

### Spillover risk increases with glacier runoff

Given these viral and eukaryotic host representations, can spillover risk be assessed in these environments? To address this question, we resorted to the novel global-fit model Random TaPas, which computes the congruence between the virus and the eukaryotic host trees, with large and weakly congruent topologies indicating low (small normalised Gini coefficient *G*^*\**^) and high (large *G*^*\**^) spillover risk, respectively.

Different patterns of spillover risks were observed in soil and in lake sediments, with both GD (figure 3) and PACo (figure S8) exhibiting consistent results. In soil, when runoff volume is negligible (C-Soil; figures 3 and S8), spillover risk is low, as median *G*^*\**^ ranges between 0.60 and 0.61, thus below the 2/3 threshold. When runoff increases from low (L-Soil) to high (H-Soil), the median *G*^*\**^ increases to 0.75, but then decreases to 0.72 for GD (figure 2), with similar values for PACo (figure S8). As such, our results show that as runoff volume increases from low to high, spillover risk remains high, but declines in soil samples (Dunn test, BH correction, *P* « 0.001). A possible explanation is that as glacial runoff increases, so does the erosion force of the glacier, which transports the content of the riverbed and riverbanks into the lake. This erosion would hence remove organisms from the topsoil this environment, and hence curb the chance of interactions between viruses and hosts, that is, limit spillover risk.

**Figure 3.**
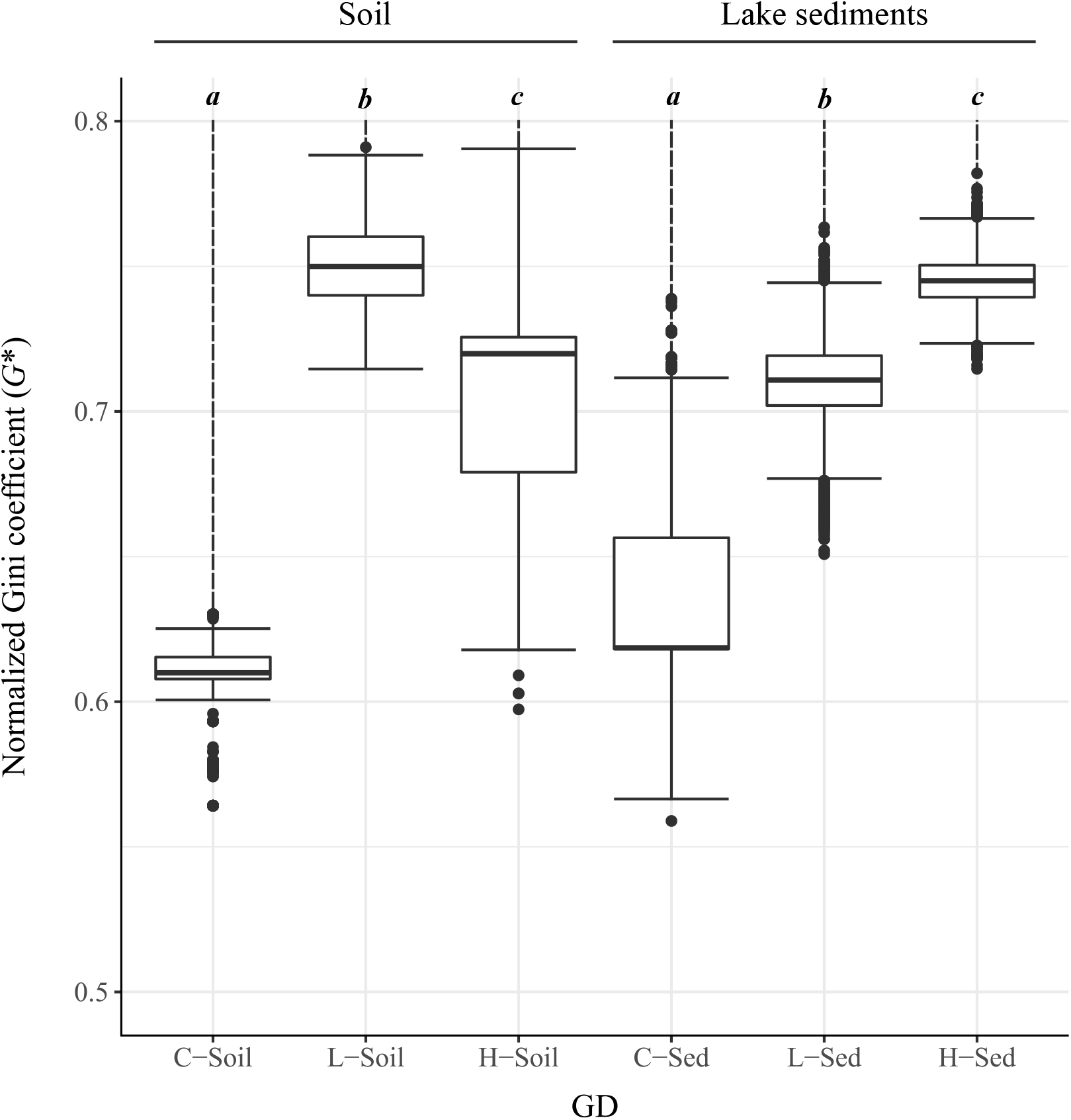
Normalised Gini coefficients (*G*^*\**^) obtained with Random TaPas (*n* = 10 runs). The values are separated by runoff volume: control (C sites), low (L sites), and high runoff (H sites). The global-fit model used was GD (geodesic distances in tree space). Significant results (Dunn test, BH correction) within each site (*n* = 10 ×1, 000_*replicates*_ for each site) are marked with letters from *a* to *c* (*α* = 0.01). Outliers are shown as dots. Similar results were found with the PACo model (figure S8).

Similarly, in lake sediments, spillover risk’s median *G*^*\**^ ∈ [0.62, 0.67] for C-Sed (figures 3 and S8), again representing low to medium spillover risk. However, as runoff increases from low to high, *G*^*\**^ increases for both GD (L-Sed: 0.71, H-Sed: 0.75; figure 3) and PACo (L-Sed 0.75, H-Sed: 0.76; figure S8), although the medians remains high for both sites. Altogether, and in contrast to soil samples, these results show that an increase in runoff volume in lake sediments significantly increases spillover risk (Dunn test, BH correction, *P* « 0.001).

This last pattern is consistent with the predictions of the Coevolution Effect hypothesis (Zohdy et al., 2019), and provides us with a mechanism explaining the observed increase in spillover risk with runoff. Lake Hazen was recently found to have undergone a dramatic change in sedimentation rates since 2007 compared to the previous 300 years: an increase in glacial runoff drives sediment delivery to the lake, leading to increased turbidity that perturbs anoxic bottom water known from the historical record (Lehnherr et al., 2018). Not only this, but turbidity also varies within the water column throughout the season (St. Pierre et al., 2019), hence fragmenting the lake habitat every year, and more so since 2007. This fragmentation of the aquatic habitat (leading up to varves in undisturbed lakes) creates conditions that are, under the Coevolution Effect, favourable to spillover. Fragmentation creates barriers to gene flow, that increases genetic drift within finite populations, accelerating the coevolution of viruses and of their hosts. This acceleration leads to viral diversification, as shown by an increasing *β*-diversity that, from the C to the H pairs of sites, goes from 21, 28, and 40 at the level of viral families (figure S7). In turn, should this diversification be combined with “bridge vectors” (such as mosquitoes in terrestrial systems) and/or invasive reservoir species, a corresponding increase in spillover risk could be expected (Zohdy et al., 2019). Lake sediments are environmental archives: over time, they can preserve genetic material from aquatic organisms but also, and probably to a lesser extent, genetic material from its drainage basin. The coevolutionary signal detected in lake sediments reflects interactions that may have happened in the fragmented aquatic habitat but also elsewhere in the drainage basin. Regardless of where the interaction occurred, our results show that spillover risk increases with runoff, a proxy of climate warming, in lake sediments (figures 3 and S8).

To our knowledge, this is the first attempt to assess the complete virosphere of both DNA and RNA viruses, and their spillover capacity in any given environment, leading us to show that increased glacier runoff, a direct consequence of climate change, is expected to increase viral spillover risk in the sediments of Lake Hazen. However, as this is the first study applying the Random TaPas algorithm, it is difficult to gauge how large *G*^*\**^ needs to be to reflect large spillover risks. Alternatives algorithms that estimate virus/host cophylogenies such as Coala (Baudet et al., 2014) generally reconcile both trees by minimising a number of events (*e*.*g*., codiversification, duplication, loss, or host switch), and hence may not offer a direct measure of spillover risk. Furthermore, our study was limited by its number of replicates, as we only had five metagenomic libraries *per* sample (*n* = 2 and *n* = 3 for DNA and RNA, respectively). Additional sampling throughout the High Arctic would also be necessary to further reinforce our results, and to calibrate the “true” risk of viral spillovers. We finally note that, because our approach relies on *known* virus/host associations, it is impossible to anticipate spillovers into novel hosts, as it was the case for viruses causing HIV (Sharp & Hahn, 2011), or through intermediate hosts as with SARS (Hu et al., 2017), and probably COVID-19 (Platto et al., 2021). As a result, our quantification of spillover risk is likely to represent a lower bound of the actual risk.

### Spillovers might already be happening

To go one step further and identify the viruses and host kingdoms that are most at risk of spillovers, we focused on the model predictions made by Random TaPas. Under the null model, the occurrence of each virus/host association is evenly distributed on their cophylogeny (when sub-cophylogenies are drawn randomly, from a uniform law). Departures from an even distribution are measured by the residuals of the linear fit. Positive residuals indicate a more frequent association than expected, that is, pairs of virus/host species that contribute the most to the cophylogenetic signal. On the other hand, negative residuals indicate a less frequent association than expected, and hence pairs of virus/host species that contribute little to the cophylogenetic signal, because they tend to create incongruent phylogenies, a signature of spillover risk (figures S9 and S10).

When examining the virus/host associations of the ten most negative residuals of each Random TaPas run (*n* = 10), all samples confounded (*n* = 6), we found that animals and protists are the most susceptible to spillover, as they exhibit the lowest median residuals for both GD (−13.08 and −20.13; figure 4) and PACo (−14.37 and −24.52; figure S11). On the other hand, plants and fungi showed a lower susceptibility to spillovers, as their median residuals were significantly higher (plants: −9.02 [GD] and −3.49 [PACo]; fungi: −3.33 [GD] and −3.95 [PACo]; all *P* « 0.01) – even though plant and fungal viruses were overrepresented in our samples (figures 4 and S11). Taking a more stringent threshold with the five most negative residuals led to similar results (figure S12).

**Figure 4.**
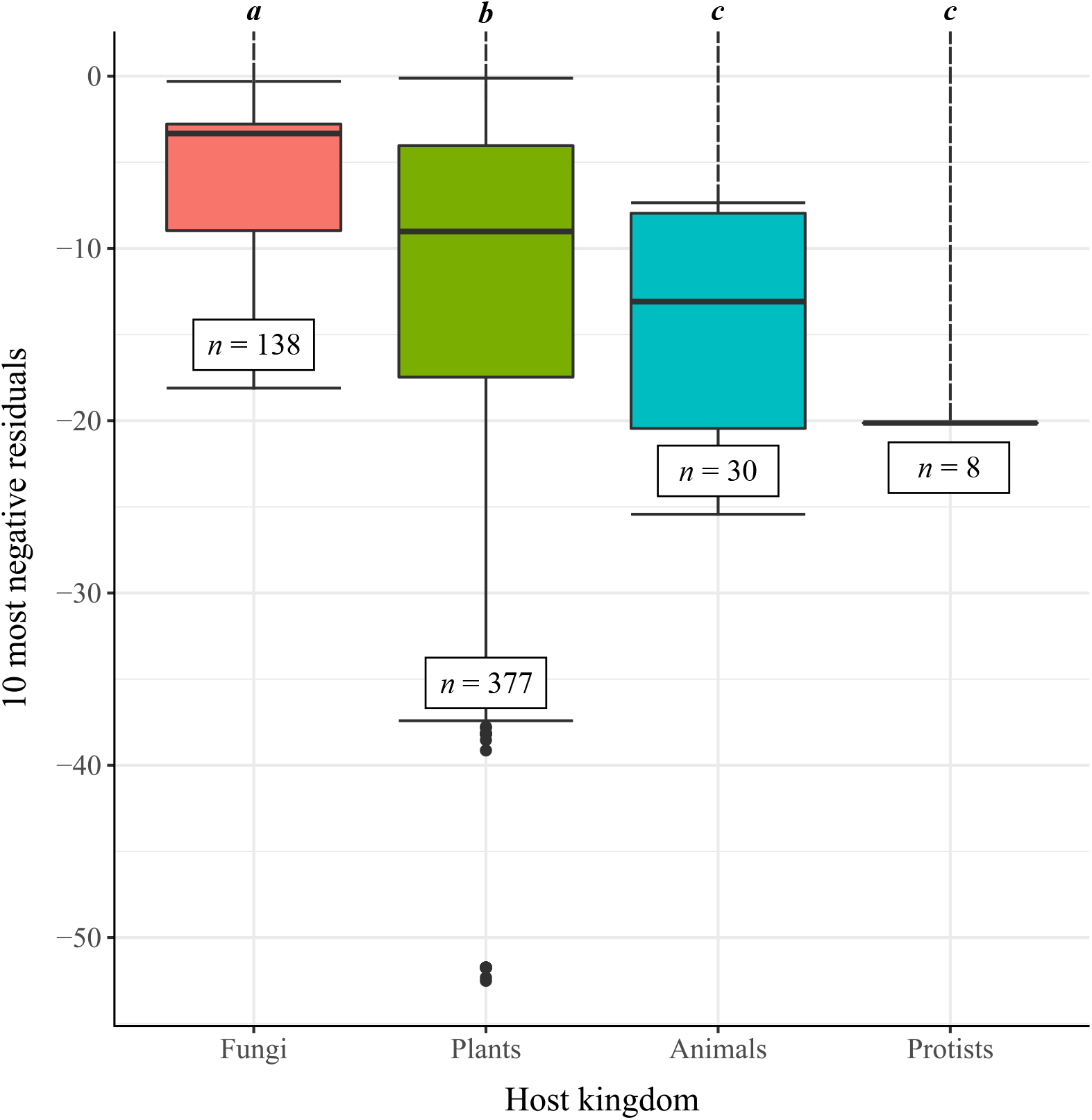
Eukaryotic hosts most susceptible to viral spillovers. The residuals were computed by Random TaPas (*n* = 10 runs) using GD (geodesic distances in tree space), all samples (*n* = 6) confounded, from which the ten most negative virus/host associations were analysed (total sample size: *n* = 10 × 6 × 10_*MostNeg*_ *− PosRes* = 553). Significant results (Dunn test, BH correction) are marked with letters from *a* to *c* (*α* = 0.01). Outliers are shown as dots. Figure S11 further shows these results to be robust to the distance used to compare trees.

To further assess the robustness of this general finding, that animals and protists are most susceptible to viral spillovers in the High Arctic, we next examined the associations with the most negative residuals in each environment separately – as we showed above that spillover risk increases in lake sediments only. We found that for both GD (figure S13*a*-*c*) and PACo (figure S13*b*-*d*), the most negative residuals were associated with animals and/or protists. Some of these hosts include the relatives of known disease vectors such as the yellow fever mosquito (*Aedes aegypti*) or the deer tick (*Ixodes scapularis*), and even pathogens such as *Pseudogymnoascus destructans*, which causes white-nose syndrome in bats, or *Fusarium poae*, involved in plant wilting (tables S7 and S8) – hence suggesting that spillovers might already be happening.

Altogether, we provided here a novel approach to assessing spillover risk. This is not the same as predicting spillovers or even pandemics, first because we rely on *known* virus/host associations, and also because as long as viruses and their “bridge vectors” are not simultaneously present in the environment (Zohdy et al., 2019), the likelihood of dramatic events probably remains low. But as climate change leads to shifts in species ranges and distributions, new associations can emerge (Wallingford et al., 2020), bringing in vectors that can mediate viral spillovers (Rocklöv & Dubrow, 2020), as simulations recently highlight (Carlson et al., 2022). This twofold effect of climate change, both increasing spillover risk and leading to a northward shift in species ranges (Osland et al., 2021), could have dramatic effect in the High Arctic. Disentangling this risk from actual spillovers and pandemics will be a critical endeavour to pursue in parallel with surveillance activities, in order to mitigate the impact of spillovers on economy and health-related aspects of human life, or on other species (Becker et al., 2019).

## Materials and Methods

### Data acquisition

An overview of data acquisition and analytical pipeline is shown in figure 1. Between the 10^th^ of May and the 10^th^ of June, 2017, sediment and soil cores were collected from Lake Hazen (82^*◦*^N, 71^*◦*^W; Quttinirpaaq National Park, northern Ellesmere Island, Nunavut, Canada), the largest High Arctic lake by volume in the world, and the largest freshwater ecosystem in the High Arctic (Lehnherr et al., 2018). Sampling took place as the lake was still completely covered in ice (table S1), as previously described (Colby et al., 2020). The sediment accumulation at the bottom of the Lake is caused by both allochthonous and autochthonous processes. The former are characterised by meltwaters that flow between late June and the end of August, and run from the outlet glaciers along the northwestern shoreline through poorly consolidated river valleys, while the latter refer to the sedimentation process within the lake.

To contrast soil and sediment sites, core samples were paired, whenever possible, between these two environments. Soil samples were taken at three locations (figure S1; C-Soil, L-Soil, and H-Soil) in the seasonally dried riverbeds of the tributaries, on the northern shore, upstream of the lake and its sediments. The corresponding paired lake sediment samples were also cored at three locations, separated into hydrological regimes by seasonal runoff volume: negligible, low, and high runoff (figure S1; C-Sed, L-Sed, and H-Sed). Specifically, the C (for *Control*) sites were both far from the direct influence of glacial inflows, while the L sites were at a variable distance from Blister Creek, a small glacial inflow. The H sites were located adjacent to several larger glacial inflows (Abbé River and Snow Goose). The water depth at L-Sed and H-Sed was respectively 50 and 21 m, and the overlying water depth for site C-Sed was 50 m.

Before sample collection, all equipment was sterilised with both 10% bleach and 90% ethanol, and non-powdered latex gloves were worn to minimise contamination. Three cores of ∼ 30 cm length were sampled at each location, and the top 5 and 10 cm of each sediment and soil core, respectively, were then collected and homogenized for genetic analysis. DNA was extracted on each core using the DNeasy PowerSoil Pro Kit, and RNA with the RNeasy PowerSoil Total RNA Kit (MO BIO Laboratories Inc, Carlsbad, CA, USA), following the kit guidelines, except that the elution volume was 30 *µ*L. DNA and RNA were thereby extracted three times per sampling site, and elution volumes were combined for a total volume of 90 *µ*L instead of 100 *µ*L.

We resorted to shotgun metagenomics and metatranscriptomics to sequence the entire DNA and RNA content of each sample, and thus the genes of all the organisms present in our environments. To do so, a total of 12 metagenomic libraries were prepared (*n* = 6 for DNA, *n* = 6 for RNA), two for each sampling site, and run on an Illumina HiSeq 2500 platform (Illumina, San Diego, CA, USA) at Génome Québec, using Illumina’s TruSeq LT adapters (forward: AGATCGGAAGAGCACACGTCTGAACTCCAGTCAC, and backward: AGATCGGAAGAGCGTCGTGTAGGGAAAGAGTGT) in a paired-end 125 bp configuration. Each library was replicated (*n* = 2 for DNA, *n* = 3 for RNA) for each sample. DNA and RNA yields following extractions can be respectively found in previous work (Colby et al., 2020) and in table S2.

### Data preprocessing and taxonomic assignments

A first quality assessment of the raw sequencing data was made using FastQC v0.11.8 (Andrews et al., 2010). Trimmomatic v0.36 (Bolger et al., 2014) was then employed to trim adapters and low-quality reads and bases using the following parameters: `phred33, ILLUMINACLIP:adapters/TruSeq3-PE-2. fa:3:26:10, LEADING:3, TRAILING:3, SLIDINGWINDOW:4:20, CROP:105, HEADCROP:15, AVGQUAL:20, MINLEN:36`. A second round of quality check was performed with FastQC to ensure that Illumina’s adapter sequences and unpaired reads were properly removed. Reads assembly into contigs was done *de novo* with both SPAdes v3.13.1 (Prjibelski et al., 2020) and metaSPAdes v3.13.1 (Nurk et al., 2017) for DNA, and with Trinity v2.9.0 (Grabherr et al., 2011), rnaSPAdes v3.13.1 (Bushmanova et al., 2019), and metaSPAdes for RNA. In all cases, the pipelines were used with their default settings. We chose metaSPAdes and rnaSPAdes for the DNA and RNA data, respectively, based on (i) the number of contigs generated, (ii) the taxonomic annotations, (iii) the time of assembly, and (iv) the contig lengths (see Electronic Supplementary Material).

Once assembled, a high-level (superkingdom) taxonomic assignment was determined based on BLASTn v2.10.0 (Z. Zhang et al., 2000) searches. Those were performed at a stringent 10^*−*19^ *E*-value threshold against the partially non-redundant nucleotide (nr/nt) database from NCBI v5 (Nucleotide [Internet]. Bethesda (MD): National Library of Medicine (US), National Center for Biotechnology Information; [1988] - [cited 2021 Jul 29]. Available from https://www.ncbi.nlm.nih.gov/nucleotide/, n.d.) (ftp.ncbi.nlm.nih.gov/blast/db/nt*tar.gz; downloaded on the 17^th^ of June, 2020). This threshold was chosen to increase the significance of our hits, as our preliminary results showed less ambiguity with smaller *E*-values, starting at a 10^*−*19^ cut-off. The proportions of taxonomic annotations (“Archaea,” “Bacteria,” “Eukaryota,” or “Viruses”) were then calculated, and a 95% consensus was taken to assign a superkingdom rank for each contig. When no such 95% consensus could be determined, the contigs were classified as “Other.” We also tested VirFinder and its trained model for predicting prokaryotic and eukaryotic viruses, but did not use it for downstream analyses as BLASTn alignments were found to be more sensitive (see Electronic Supplementary Material).

To refine the viral taxonomic assignments, we further assessed three alternative approaches. First, GenBank’s viral nucleotide sequences v238.0 (Sayers et al., 2019) were retrieved (ftp.ncbi.nlm.nih.gov/genbank/gbvrl*seq.gz; downloaded on the 23^rd^ of July, 2020), concatenated, converted to FASTA, and formatted for BLAST with the `makeblastdb` command. BLASTn searches against this viral database were then conducted on the viral contigs, at the same stringent 10^*−*19^ *E*-value threshold. Second, the same viral contigs were mapped against the same database, but with Bowtie2 v2.3.5.1 (Langmead & Salzberg, 2012), using default settings. Third, MetaGeneMark v3.38 (Besemer & Borodovsky, 1999; Zhu et al., 2010), which implements a hidden Markov model (HMM) used to predict genes of prokaryotic and eukaryotic viruses (Nishimura et al., 2017; Laffy et al., 2016; Gupta et al., 2016), was used to predict protein-coding regions in the viral contigs, followed by BLASTp v2.12.0 searches against RefSeq’s v210 (O’Leary et al., 2015) viral protein database (ftp.ncbi.nlm.nih.gov/refseq/release/viral/*faa.gz; downloaded on the 15^th^ of February, 2022). As we found that the MGM + BLASTp approach was the most sensitive (a 4× to 15× increase; see table S6), this is what results from this point on are based on.

To refine the eukaryotic taxonomic assignments, we followed a similar approach. As GeneMark only implements five eukaryotic models, we employed Augustus v3.4.0 (Stanke et al., 2008) to predict the eukaryotic protein-coding regions. This HMM implements 109 models available, including 105 eukaryotic species, spanning 76 genera. To avoid using similar models multiple times, we randomly chose one species *per* genus, attempting to cover the entire superkingdom. Predictions were filtered at a *P*-value threshold of 0.05, and the eukaryotic model with the smallest *P*-value was then kept. In case of ties, the corresponding contigs were removed from the analyses. BLASTp searches were conducted as described above, on the following six RefSeq’s protein divisions (downloaded on the 15^th^ of February, 2022): fungi, invertebrate, plant, protozoa, vertebrate (“mammalian”), and vertebrate (“other”).

For each sampling location, after combining the genes predicted from both the DNA and RNA contigs classified as “viral” and “eukaryotic,” the first 12 High-scoring Segment Pairs (HSPs) of each gene were kept. Those were then filtered with an *E*-value threshold of 10^*−*50^. The accession numbers of these hits were used to retrieve their corresponding taxonomy identifiers and their full taxonomic lineages with the R packages `rentrez` v1.2.3 (Winter, 2017) and `taxonomizr` v0.5.3 (Sherrill-Mix, 2019). Bacteriophages and virophages data were filtered out.

We retrieved the phylogenetic placements of all taxonomic assignments from the Tree of Life (ToL) (tolweb.org), hence generating two trees: one for the viruses and one for the eukaryotes that we identified, down to the species level. For this, we used the `classification` and `class2tree` functions from the R package `taxize` v0.9.99 (S. Chamberlain et al., 2020; S. A. Chamberlain & Szocs, 2013). At each site, vertices of the viral and eukaryotic trees were then put in relation with each other according to the Virus-Host DB (downloaded on the 24^th^ of March, 2022) (Mihara et al., 2016). These relations were saved as a binary association matrix (0: no infection; 1: infection), one for each site (see figures S5, S9-S10). The *β*-diversity, or variation in species composition, between sites *S*_1_ and *S*_2_ was computed as 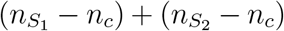 where 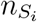 is the total number of species in site *S*_*i*_, and *n*_*c*_, the number of species that the two sites have in common.

### Spillover quantification

To quantify viral spillovers based both on the viruses and eukaryotes identified and their known associations, we employed the Random Tanglegram Partitions algorithm (Random TaPas) (Balbuena et al., 2020). This algorithm computes the cophylogenetic signal or congruence between two phylogenetic trees, the viral and the eukaryotic host trees computed as detailed above, with the normalised Gini coefficient (*G*^*\**^) – a quantitative proxy for spillover risk. Indeed, when congruence is large, or “perfect,” the two trees are identical and hence, there is strong cophylogenetic signal – and absence of spillover. On the other hand, weak congruence is evidence for the existence of spillovers. Random TaPas quantifies congruence in two ways: a geodesic distance (GD) (Schardl et al., 2008), or a Procrustes distance (Procrustes Approach to Cophylogeny: PACo) (Balbuena et al., 2013), the latter measuring the distance between two trees geometrically transformed to make them as identical as possible. To partially account for phylogenetic non-independence when measuring congruence, Random TaPas further implements a resampling scheme where *N* = 10^4^ subtrees of about 20% of the unique virus/hosts associations are randomly selected. This selection is used to generate a distribution of the empirical frequency of each association, measured by either GD or PACo.

Each empirical frequency is then regressed against a uniform distribution, and the residuals are used in two ways: (i) to quantify co-speciation, which is inversely proportional to spillover risk; and (ii) to identify those virus/host pairs that contributed the least to the cophylogenetic signal, *i*.*e*., the most to spillover risk. This risk is finally quantified by the shape of the distribution of residuals (for GD or PACo), with the normalised Gini coefficient *G*^*\**^ taking its values between 0 (perfect congruence, no spillover) to 1 (maximal spillover risk), with a defined threshold of 2/3 indicating a “large” value of *G*^*\**^ or large incongruence. To account for phylogenetic uncertainty, the process is repeated *n* = 1, 000 times, each replicate being a random resolution of the multifurcating virus/host trees of life into a fully bifurcating tree. The stability of these results was assessed by running this algorithm *n* = 10 times, and by combining the results *G*^*\**^. Post-hoc comparisons were based on the Dunn test, controlling the False Discovery Rates with the BenjaminiHochberg (BH) correction.

## Supporting information

ESM

## Data availability

The raw data used in this study can be found at www.ncbi.nlm.nih.gov/bioproject/556841 (DNA-Seq) and at www.ncbi.nlm.nih.gov/bioproject/PRJNA746497/ (RNA-Seq). The code developed for this work is available from www.github.com/sarisbro/data/, in the archive Lemieux.etal.tar.bz2.

## Acknowledgements

We thank Frances Pick for helpful comments on an early version of this work. This study was financially supported by the Natural Sciences Research Council of Canada (A.J.P., S.A.B.) and by the University of Ottawa (A.L., G.A.C.).

## Author contributions

S.A.B. and A.J.P. designed research; G.A.C. collected and processed the samples; A.L. performed all analyses and wrote the original draft; A.L. and S.A.B. wrote the manuscript with contributions and suggestions from G.A.C. and A.J.P.; and S.A.B. and A.J.P. supervised this study and acquired funding.

## Competing interests

The authors declare no competing interests.

## Notes

https://github.com/sarisbro/data

### Competing Interest Statement

The authors have declared no competing interest.

