## Supplementary material for "Viral spillover risk increases with climate change in High Arctic lake sediments": ESM

**Optimal assemblers for our data: rnaSPAdes and metaSPAdes.** Data assembly is an important step in metagenomics assessments. Some authors found that rnaSPAdes and Trinity are among the highest performing assembly tools tested [1], while others found SPAdes [2] to be performing best. To characterise viromes, both MEGAHIT and metaSPAdes [3, 4] have been recommended. Overall, previous studies showed (i) a lack of consensus in choosing an assembly tool, as results based on the same pipelines varied amongst publications, and largely depended on the available resources, the research questions, and the datasets [2]; and (ii) most studies recommended testing and comparing different tools [1].

To select an assembly pipeline to analyse our DNA data, SPAdes and metaSPAdes were tested on a single replicate of the L-Soil sample, each pipeline assembling respectively 846,341 and 1,686,159 contigs in 195.90 and 17.51 hours, respectively. While their overall distributions of contig lengths were similar, SPAdes filtered out contigs more aggressively than metaSPAdes did, their N50 being respectively 302 and 257 (figure S2), meaning that the lengths of half of the contigs were at least of 302 (SPAdes) and 257 bp (metaSPAdes). As for the taxonomic annotations, BLASTn classified 42.75% of SPAdes' contigs, while it was only 35.98% for metaSPAdes (table S3). The bacteria and viruses were respectively the most and the least abundant superkingdoms found in the DNA sample (SPAdes: 96.34% and 0.01%; metaSPAdes: 97.06% and 0.00%). Considering that metaSPAdes generated 1.99 times more contigs than SPAdes and was 11.19 times faster, we decided to assemble the DNA contigs with metaSPAdes.

Trinity, rnaSPAdes, and metaSPAdes were tested on an RNA replicate of the L-Soil sample. Assembled reads led to 727,160, 1,564,307, and 672,527 contigs for Trinity, rnaSPAdes, and metaSPAdes respectively, and required 221.59, 4.47, and 4.76 hours to complete. Trinity filtered out contigs more aggressively than rnaSPAdes, hence artificially inflating its N50 from 169 to 289 by eliminating the shortest contigs, while keeping the overall distributions of contig lengths very similar (figure S3). However, unlike Trinity, metaSPAdes did not filter out the shortest contigs, despite its N50 being 242. The shortest contigs assembled with metaSPAdes were approximatively of length 50 bp, while it was 100 bp for rnaSPAdes.

One notable difference between the three assemblies was about the taxonomic assignments: 56.44% and 49.16% of the contigs assembled with Trinity and metaSPAdes were given a taxonomy annotation, while it was only 35.74% for rnaSPAdes (table S4). The proportions of the superkingdoms "Archaea," "Eukaryota," and "Viruses" were negligible for the replicate of the L-Soil sample, as "Bacteria" made up more than 95% of all taxonomic assignments for Trinity (95.79%), rnaSPAdes (95.69%), and metaSPAdes (96.15%). The number of viral families was in fact the lowest, the assembly pipelines only identifying 177 (0.04%), 220 (0.04%), and 154 (0.05%) contigs as viral, respectively. Given that rnaSPAdes and metaSPAdes were 49.57 and 46.55 faster than Trinity, that rnaSPAdes generated the largest number of contigs, and that Trinity's and rnaSPAdes' overall lengths distributions were similar and easier to explain than that of metaSPAdes, we decided to assemble the RNA contigs with rnaSPAdes.

Differences in the number of contigs generated, the time of assembly, and the contig lengths were noted between Trinity, rnaSPAdes, and metaSPAdes. Our results therefore highlight the importance of testing multiple assembly tools to select the best-fit for our metagenomics data, as its choice can affect downstream analysis such as taxonomic assignments.

**Assigning superkingdoms to contigs using BLASTn.** DNA and RNA contigs were assigned taxonomic annotations at the superkingdom level with a similarity approach based on BLASTn and a 95% consensus. This was compared to VirFinder v1.1 [5], which relies on a machine learning algorithm that uses $k$-mer distributions rather than similarity to identify viral contigs. To contrast the number of viral contigs classified by each approach, we first downloaded VirFinder's pre-trained model to predict prokaryotic and eukaryotic genes (`https://github.com/jessieren/VirFinder/raw/master/EPV/VF.modEPV_k8.rda`; updated on the $10^{\text{th}}$ of June, 2018), and ran the algorithm on the contigs of a replicate of the L-Soil sample. In order to maximise specificity with VirFinder, we classified contigs as "viral" if the resulting $q$-value was equal to 0 (*i.e.*, $q \leq 10^{-99}$). VirFinder identified 1,625 viral contigs, while we had identified 220 with BLASTn for this specific dataset (table S4). However, only two contigs (figure S4$a$) were common to both approaches: post-hoc BLASTn analyses on VirFinder's contigs revealed that no High-scoring Segment Pairs (HSPs) in the `nt` database were found for 1,349 (83.0%) of those contigs (figure S4$b$). Furthermore, while we expected the longest contigs to have the smallest $q$-values, VirFinder mostly predicted the shortest contigs to be viral (figure S4$c$), hence rising doubts on its accuracy. As a result, and since VirFinder cannot assign contigs beyond the superkingdom level, and that our goal was to identify viral *species*, we decided to keep BLASTn annotations for the remaining steps. To apply the $\log_{10}$ transformation (figure S4$c$), note here that $q$-values $\leq 10^{-10}$ were changed to $10^{-10}$.

**Known viruses are the least dominant superkingdoms in soil and lake sediments.** To classify contigs at the superkingdom level, consensus annotations at the 95% threshold were extracted from the BLASTn alignments. Those with no assignments represented $> 50\%$ of all contigs (table S5), which is not unusual as Rosario et al. [6], for instance, found that up to 57% of their contigs returned no hits, based on a similar procedure to ours. They argued that these high proportions reflected a large fraction of unknown viruses in reclaimed water. Discovering novel viruses still remains a challenge as a similarity-based approach (*e.g.*, homology searches against current databases) cannot efficiently characterise unknown viruses [7, 8]. Thus, even if the proportions of contigs classified as "viral" were small, novel RNA and DNA viruses might be abundant in all of our samples. While these "unknown viruses" deserve more attention, this goes beyond the scope of the current study.

Out of the DNA and RNA contigs that could be annotated, most were found to be of bacterial origin ($> 72.6\%$; figure S6; table S5), while viruses represented less than 1%. This is line with previous work that either reported bacteria [9], or eukaryotes as the most abundant superkingdom [10], viruses being systematically the least abundant or not found. Note that those studies were based on amplicon sequencing (either 16S or 18S rRNA or even rDNA genes) instead of shotgun metagenomics, which might explain why viruses were mostly "missed," as no universal sequence exists for viruses, thus limiting the identification of all the viruses in an environment [11]. Yet, studies resorting to shotgun sequencing did not characterise more viruses [12, 13]. Although Metagenomics Assembled Genomes (MAGs) are now being reconstructed for viruses, this is mostly limited to bacteriophages [14], and should hence be extended to other viruses, in particular to RNA viruses, whose spillovers can be devastating [15].

**Refining the viral species annotations with MetaGeneMark (MGM) and BLASTp.** BLASTn and Bowtie2 were initially both used to refine the taxonomic annotations of the viruses. In all cases, more HSPs were found with BLASTn than with Bowtie2 (table S6), which is not unexpected as Bowtie2 expects near-exact matches between the queries and sequences in the reference database. Since we were looking for similar sequences in more than one species, hence privileging sensitivity over specificity, we only resorted to BLASTn searches. However, we note that the high sensitivity of our annotation pipeline came at the cost of low specificity, as some annotations pointed to cheetahs (*Acinonyx jubatus*), southern white rhinoceros (*Ceratotherium simum simum*), or leopards (*Panthera pardus*), which are unlikely to be found in High Arctic samples, while the human sequences might either represent actual human, indigenous, or other presence on Ellesmere Island.

As similarity searches were directly performed on the short DNA and RNA contigs, of median sizes 263 and 173 bp, respectively, which may lead to sensitivity issues, we then tried to predict protein-coding regions for the viral contigs. We resorted to the HMMs implemented in MetaGeneMark v3.38 [MGM; 16–18], followed by BLASTp searches against RefSeq's protein viral database. In all cases, the number of hits with the MGM + BLASTp combined approach surpassed those of BLASTn and Bowtie2, $>$ 1000 HSPs being returned in each sample (table S6). Considering these results, we opted for a protein-coding prediction approach (MGM + BLASTp) instead of inferring the viral species directly from the contigs (BLASTn).

Among all the viral HSPs, we found species of bacteriophages, eukaryotic viruses, and even of one virophage (table 1, figure S7). In 4 out of 6 samples, we found more viruses infecting bacteria than eukaryotes, although it was more noticeable in lake sediments (table 1), and some families of DNA viruses such as *Podoviridae*, *Myoviridae*, *Siphoviridae*, all bacteriophages, were found to be amongst the most abundant, which is consistent with other surveys from soil [19] or extreme environments [20]. In fact, these results further reinforce our methodology (*i.e.,* predicting protein-coding regions with MGM), as no bacteriophages had been found when we directly inferred the viral species from the contigs with BLASTn, which is largely inconsistent with studies previously published [20, 21].

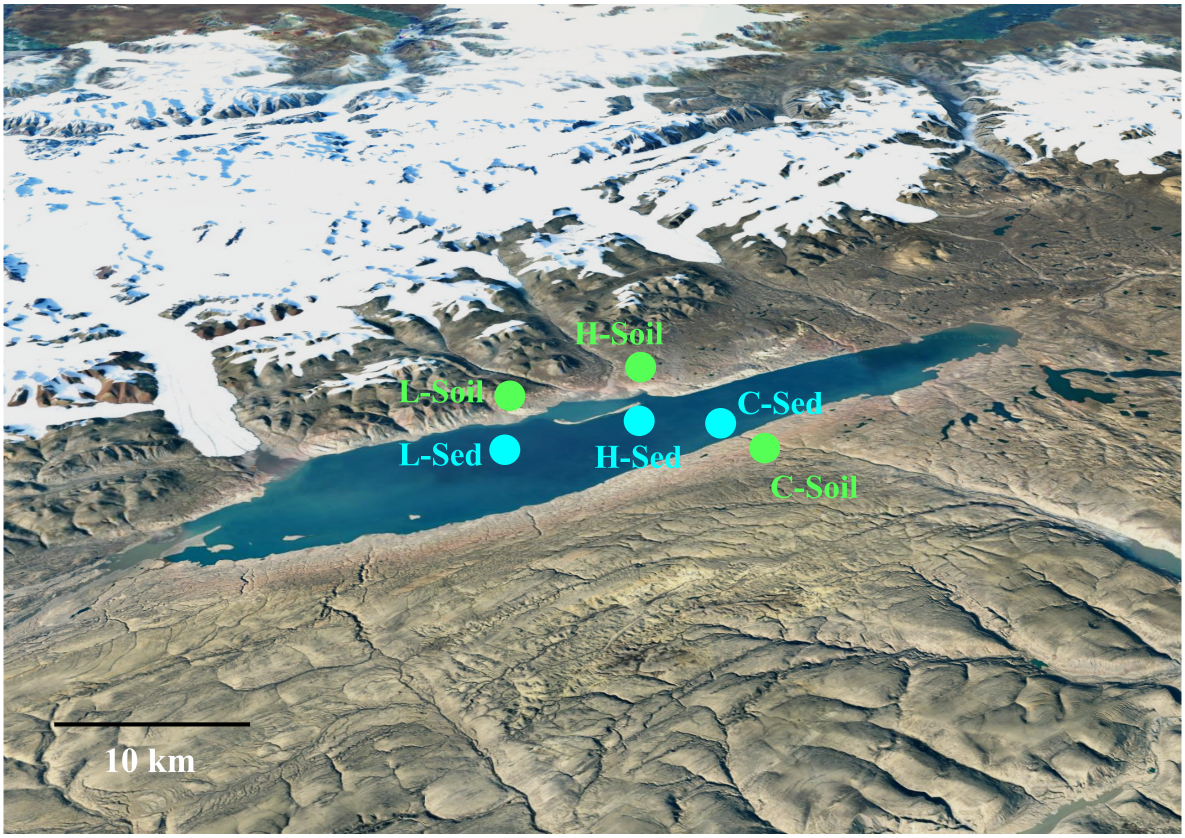


**Figure S1.** Location of Lake Hazen and sampling sites. Green and blue dots represent soil and lake sediments samples, respectively, and were separated into three hydrological regimes: negligible (C for *Control*), low (L), and high (H) runoff volume. This figure was adapted from previous work [22], and used Google Earth for terrain view.

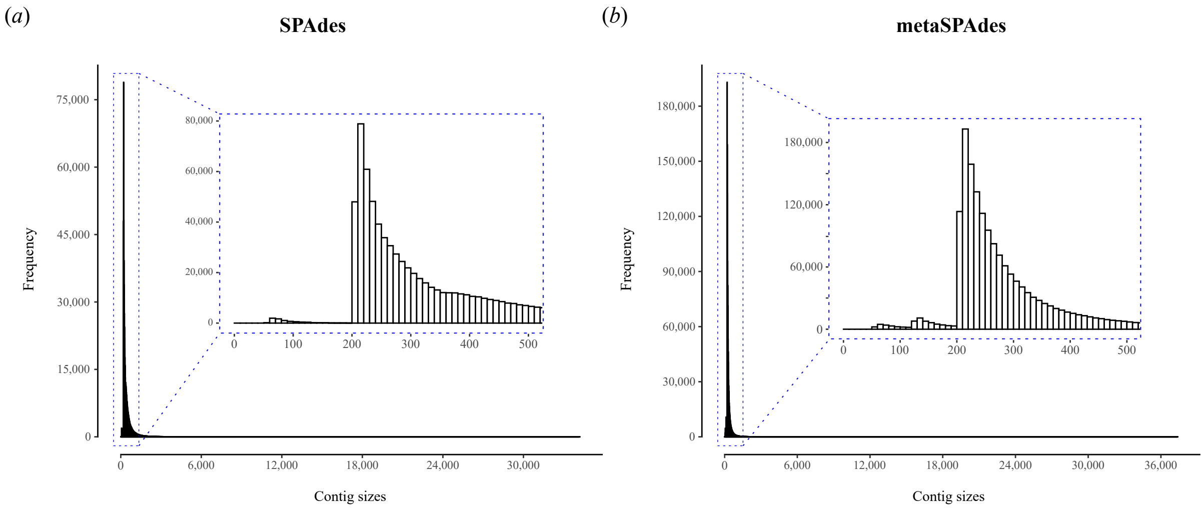


**Figure S2.** Length distribution of the DNA contigs assembled with (*a*) SPAdes and (*b*) metaSPAdes. Each assembler was assessed on a single replicate (of the L-Soil sample), leading to 846,341 and 1,686,159 contigs, respectively. The N50 statistics for SPAdes and metaSPAdes were 302 and 257.

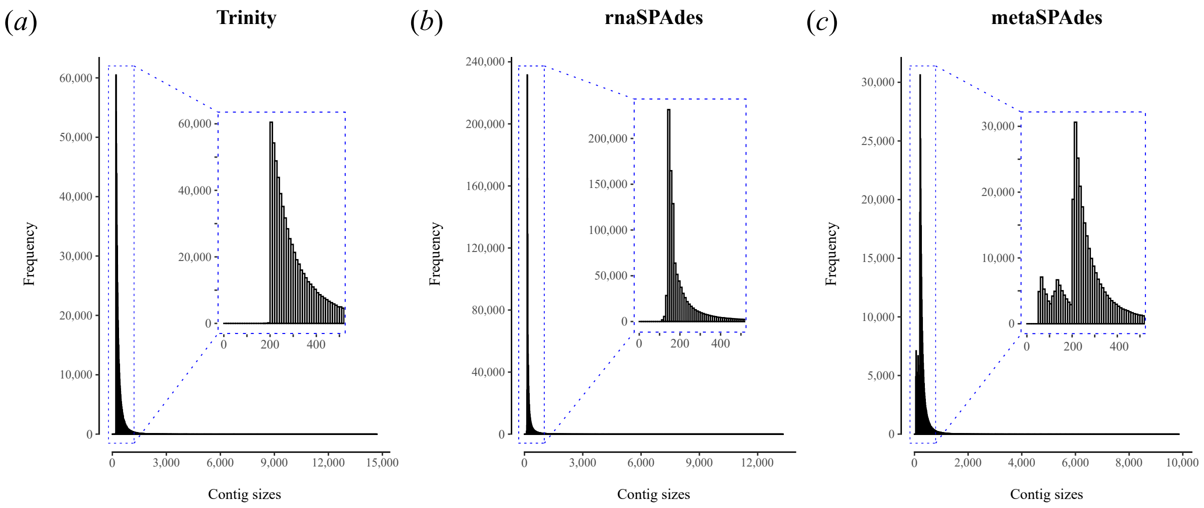


**Figure S3.** Length distribution of the RNA contigs assembled with (*a*) Trinity, (*b*) rnaSPAdes, and (*c*) metaSPAdes. Each assembler was assessed on a single replicate (of the L-Soil sample), leading to 727,160, 1,564,307, and 672,527 contigs, respectively. The N50 statistics for Trinity, rnaSPAdes, and metaSPAdes were 289, 169, and 242.

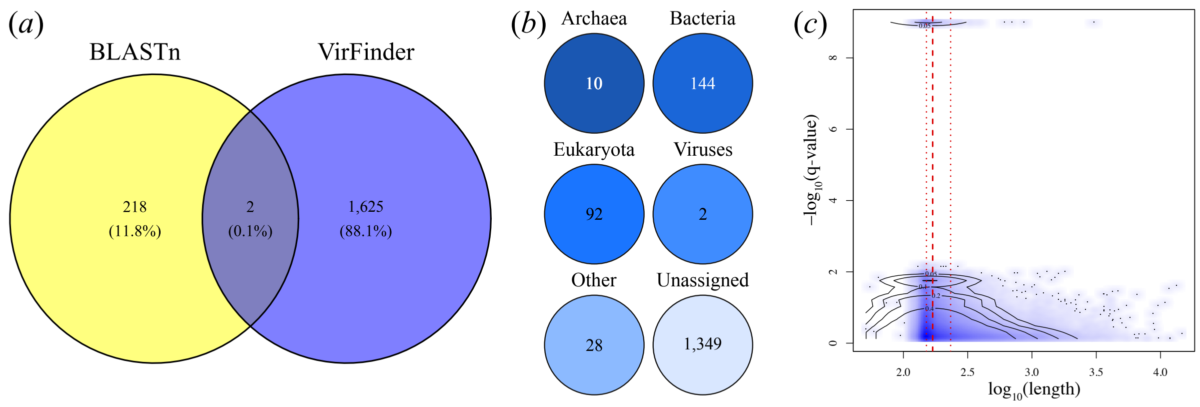


**Figure S4.** Comparison between BLASTn and VirFinder's identification of viral contigs. (*a*) Contigs identified as viral by both algorithms. (*b*) Post-hoc BLASTn annotations of the contigs identified as "viral" by VirFinder. (*c*) Distribution of VirFinder's *q*-values. The lines of quartiles (Q1, median, and Q3) are also shown.

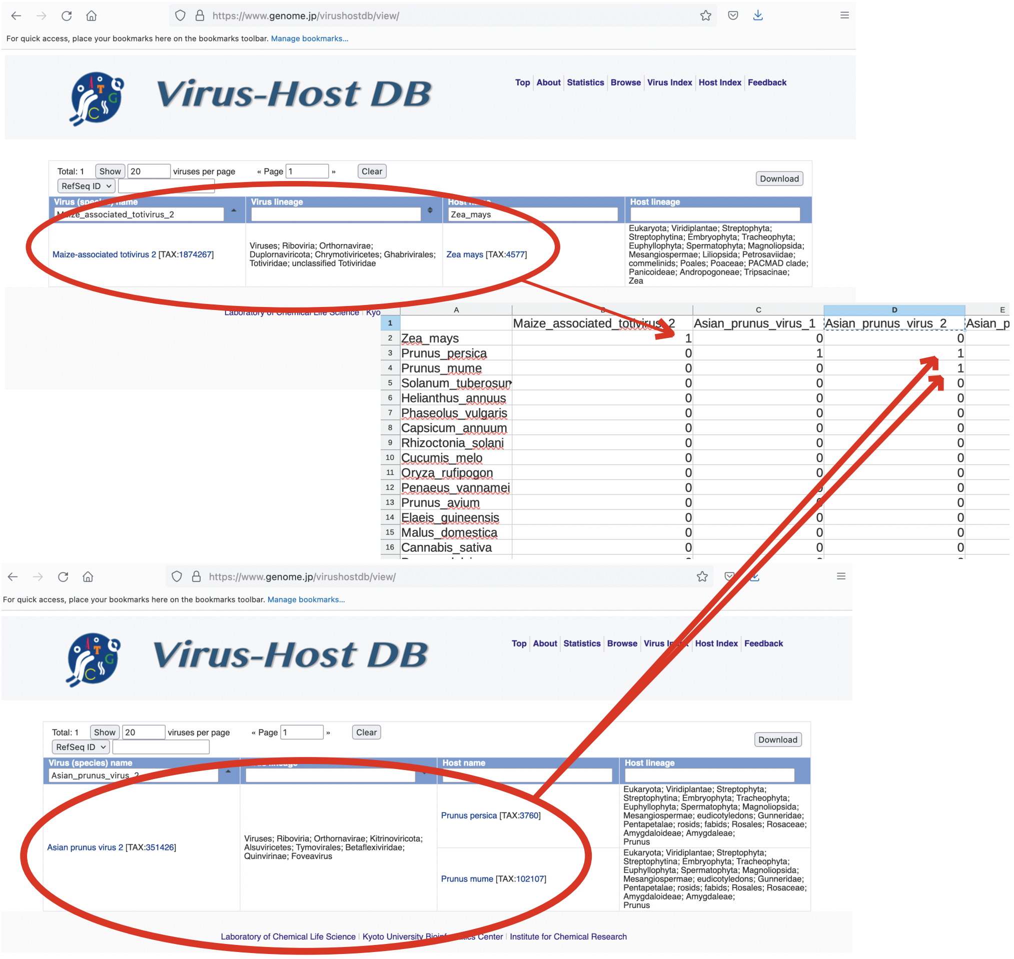


**Figure S5.** Schematic depicting how an association matrix is constructed from the Virus-Host DB. Only one host is found in the database for the *Maize-associated totivirus 2*, *Zea mays* (top), and this association is indicated by a 1, while two hosts are found for *Asian prunus virus 2*, *Prunus persica* and *Prunus mume* (bottom), and both associations are indicated by a 1. Absence of associations is indicated by 0. See `www.github.com/sarisbro/data` for the relevant script.

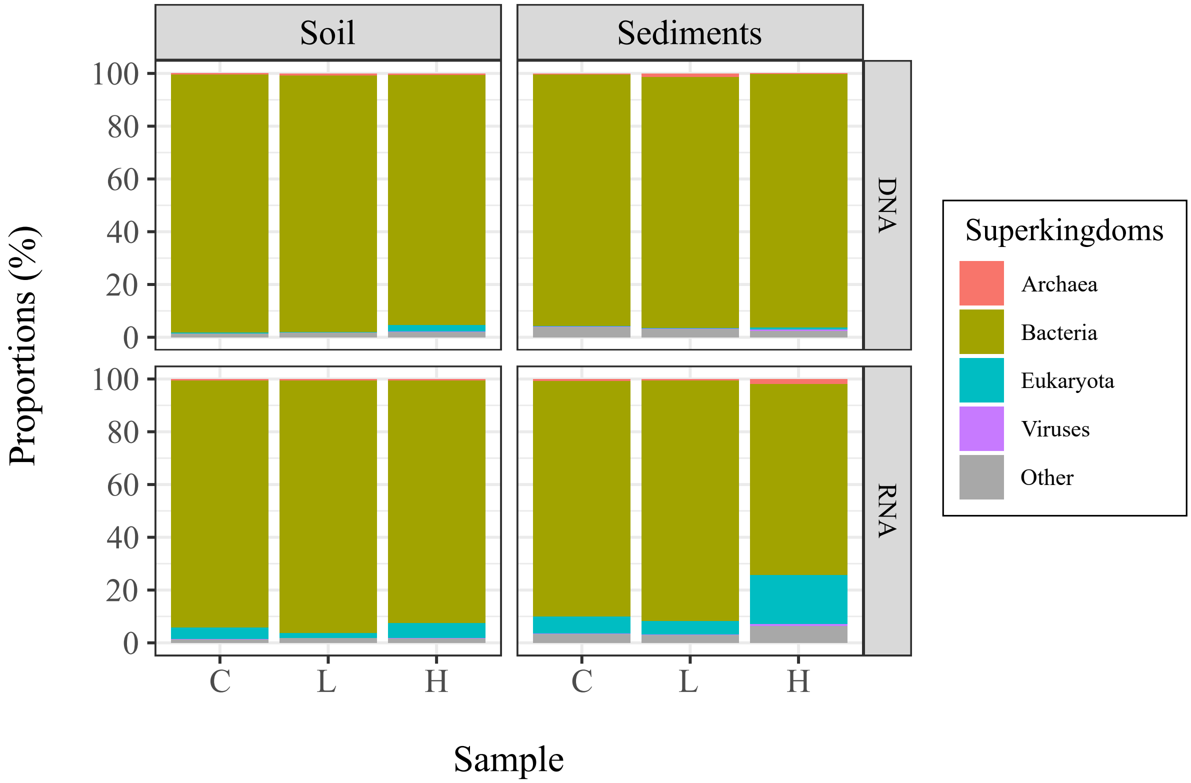


**Figure S6.** The average proportions of taxonomic annotations in soil and lake sediments. The proportions of superkingdoms were calculated for each DNA ($n = 2$ *per* sample) and RNA ($n = 3$ *per* sample) replicate, and averages were then computed per sample based on these values. The data used for this figure can be found in table S5.

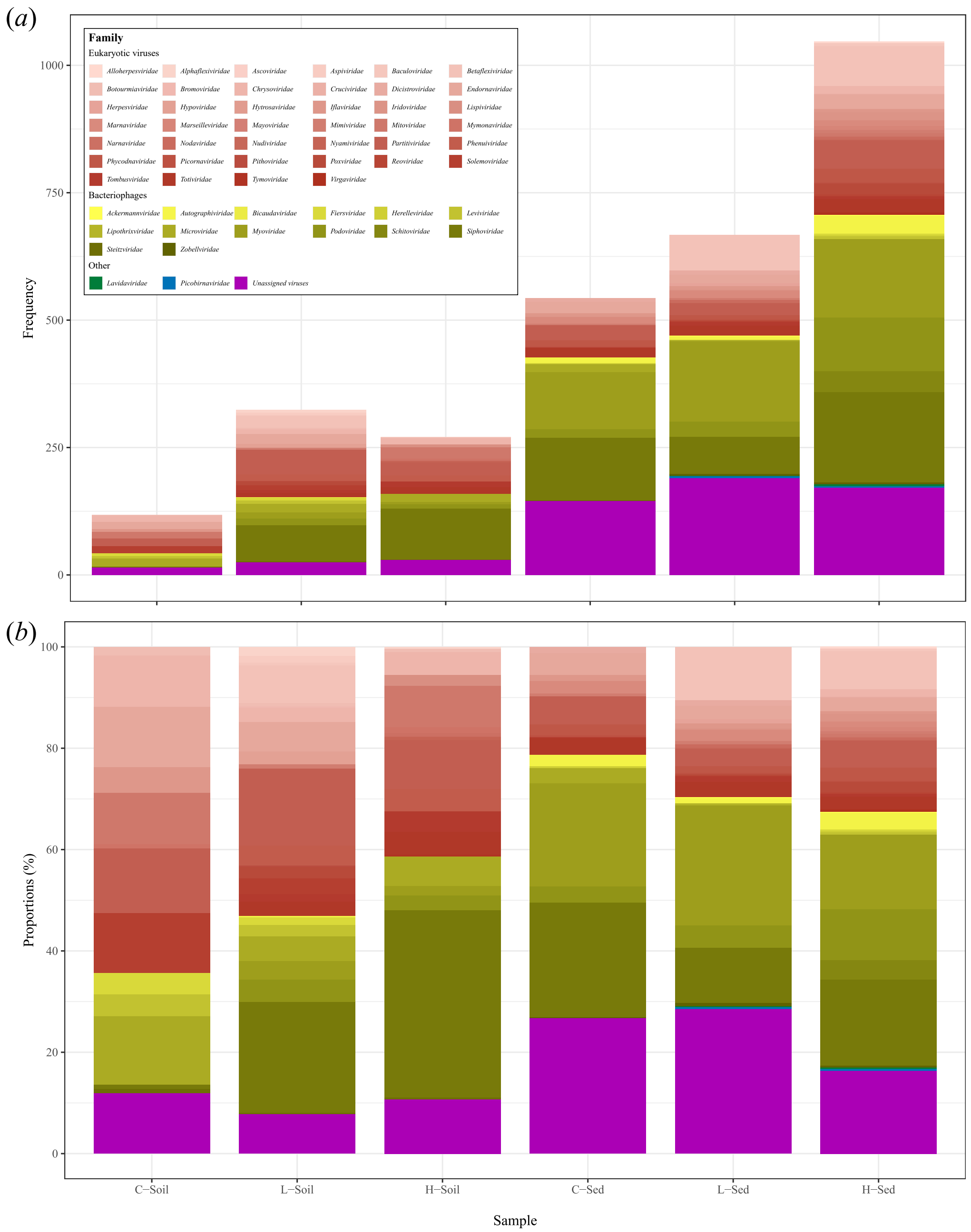


**Figure S7.** Distribution of the High-scoring Segment Pairs (HSPs) of all viruses, at the family rank. The (*a*) raw counts and the (*b*) proportions are represented. Each viral species was kept once. The data used for this figure can be found in table 1, and figure 2 shows the distribution of the eukaryotic viral families only.

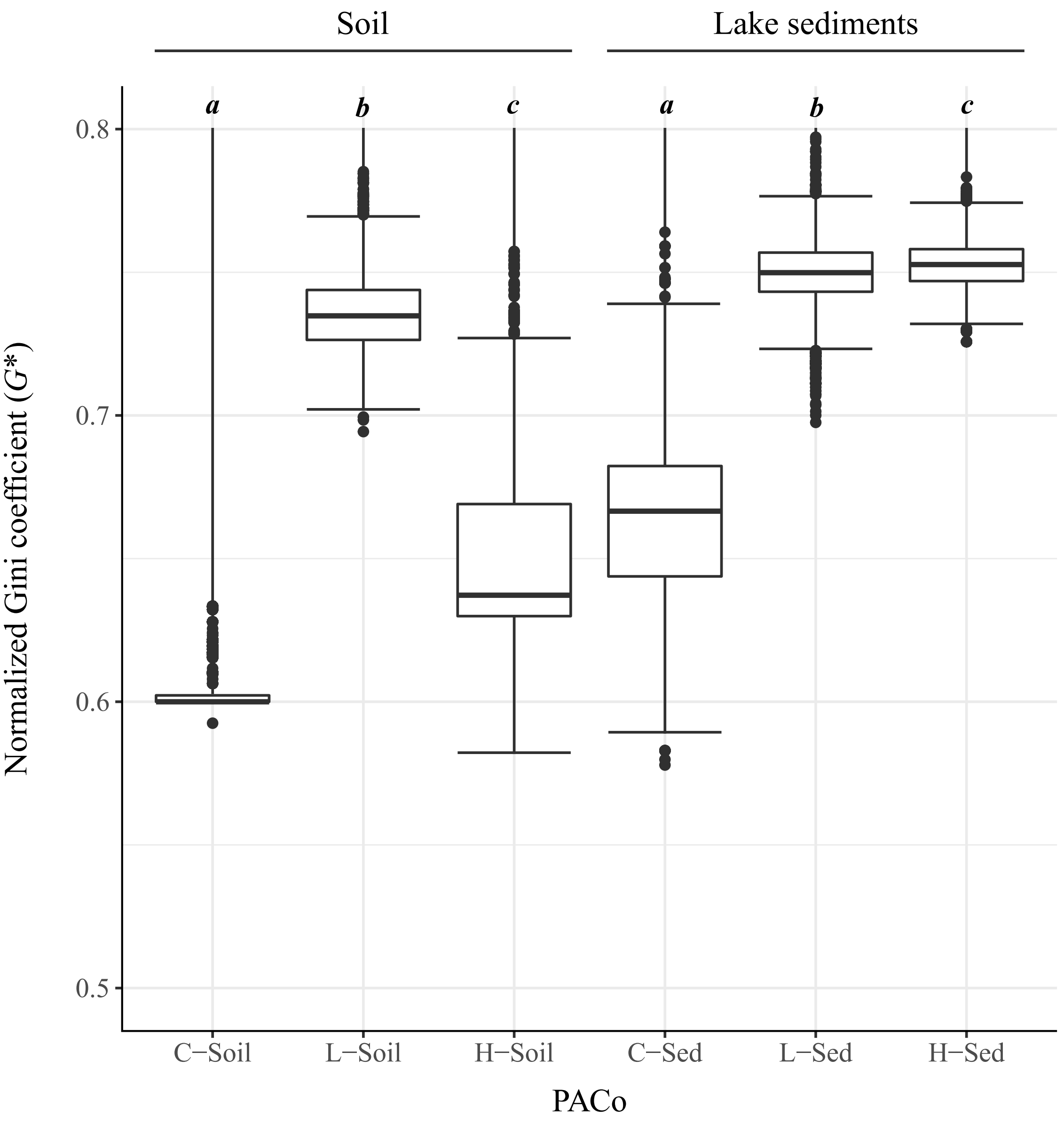


**Figure S8.** Normalised Gini coefficients ($G^{\star}$) obtained with Random TaPas ($n = 10$ runs). The values are separated by runoff volume: control (C sites), low (L sites), and high runoff (H sites). The global-fit model used was PACo (Procrustes Approach to Cophylogeny). Significant results (Dunn test, BH correction) within each site ($n = 10 \times 1,000_{replicates}$ for each site) are marked with letters from $a$ to $c$ ($\alpha = 0.01$). Outliers are shown as dots. Similar results were previously found with the GD model (geodesic distances in tree space; figure 3).

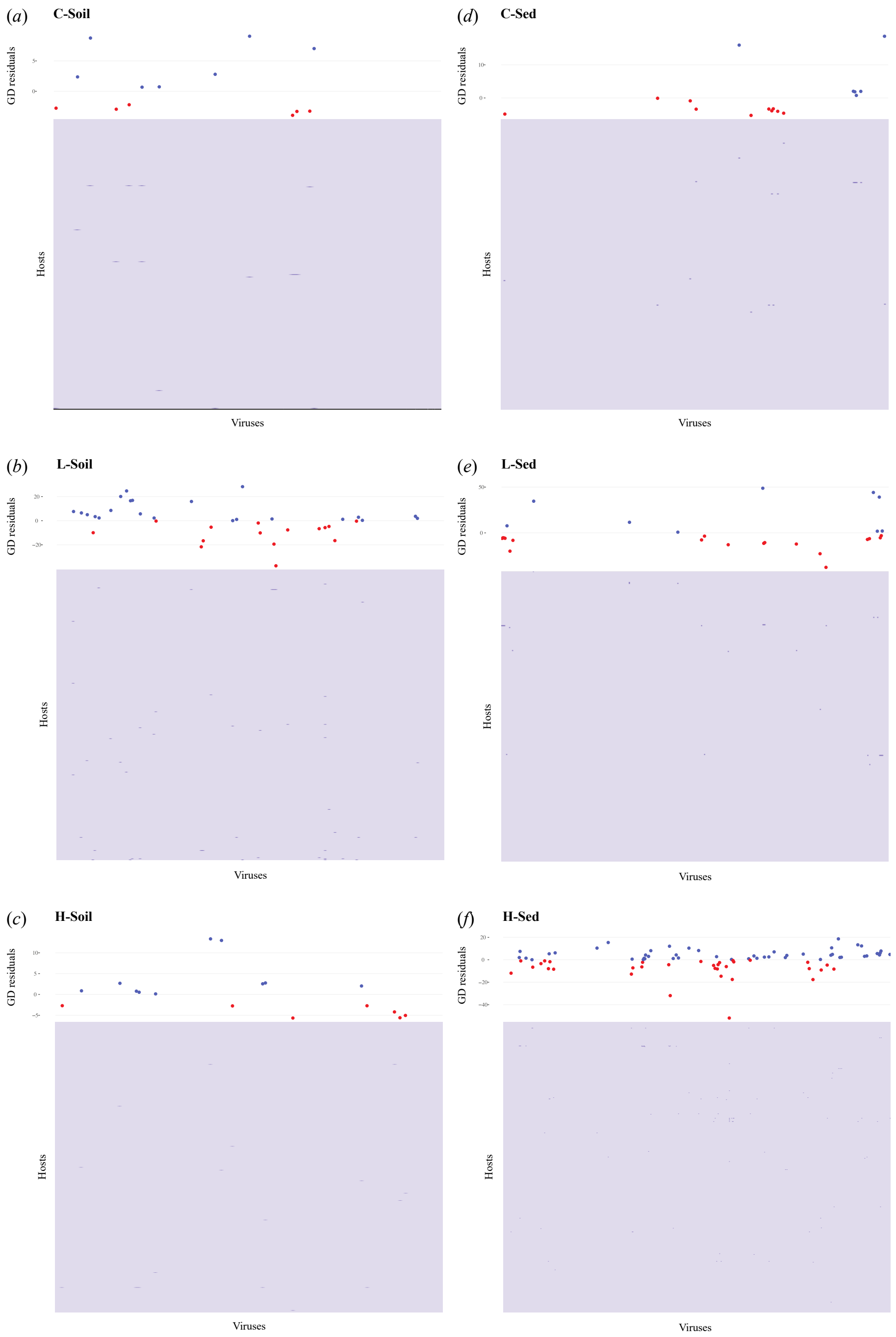


**Figure S9.** Virus/host association heatmaps and associated mean residuals, *per* sampling site. (*a*) C-Soil; (*b*) L-Soil; (*c*) H-Soil; (*d*) C-Sed; (*e*) L-Sed; and (*f*) H-Sed sites. Residuals were computed by Random TaPas ($n = 10$ runs) using GD (geodesic distances in tree space). Blue and red colours represent positive and negative residuals, respectively. In cases of viruses infecting $> 1$ host, mean residuals were calculated on all the associations.

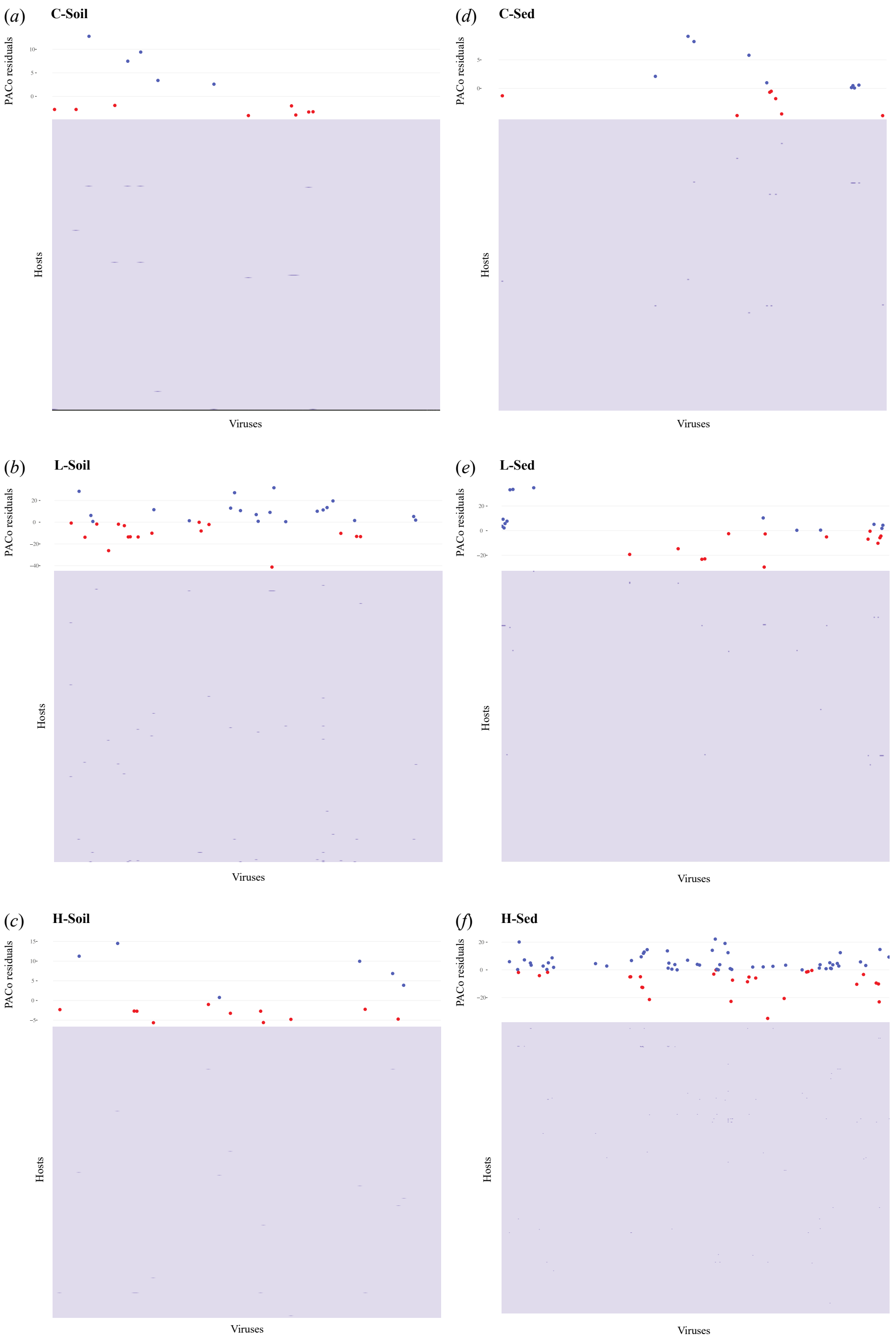


**Figure S10.** Virus-host associations heatmaps and associated mean residuals, *per* sampling site. (*a*) C-Soil; (*b*) L-Soil; (*c*) H-Soil; (*d*) C-Sed; (*e*) L-Sed; and (*f*) H-Sed sites. Residuals were computed by Random TaPas ($n = 10$ runs) using PACo (Procrustes Approach to Cophylogeny). Blue and red colours represent positive and negative residuals, respectively. In cases of viruses infecting $> 1$ host, mean residuals were calculated on all the associations.

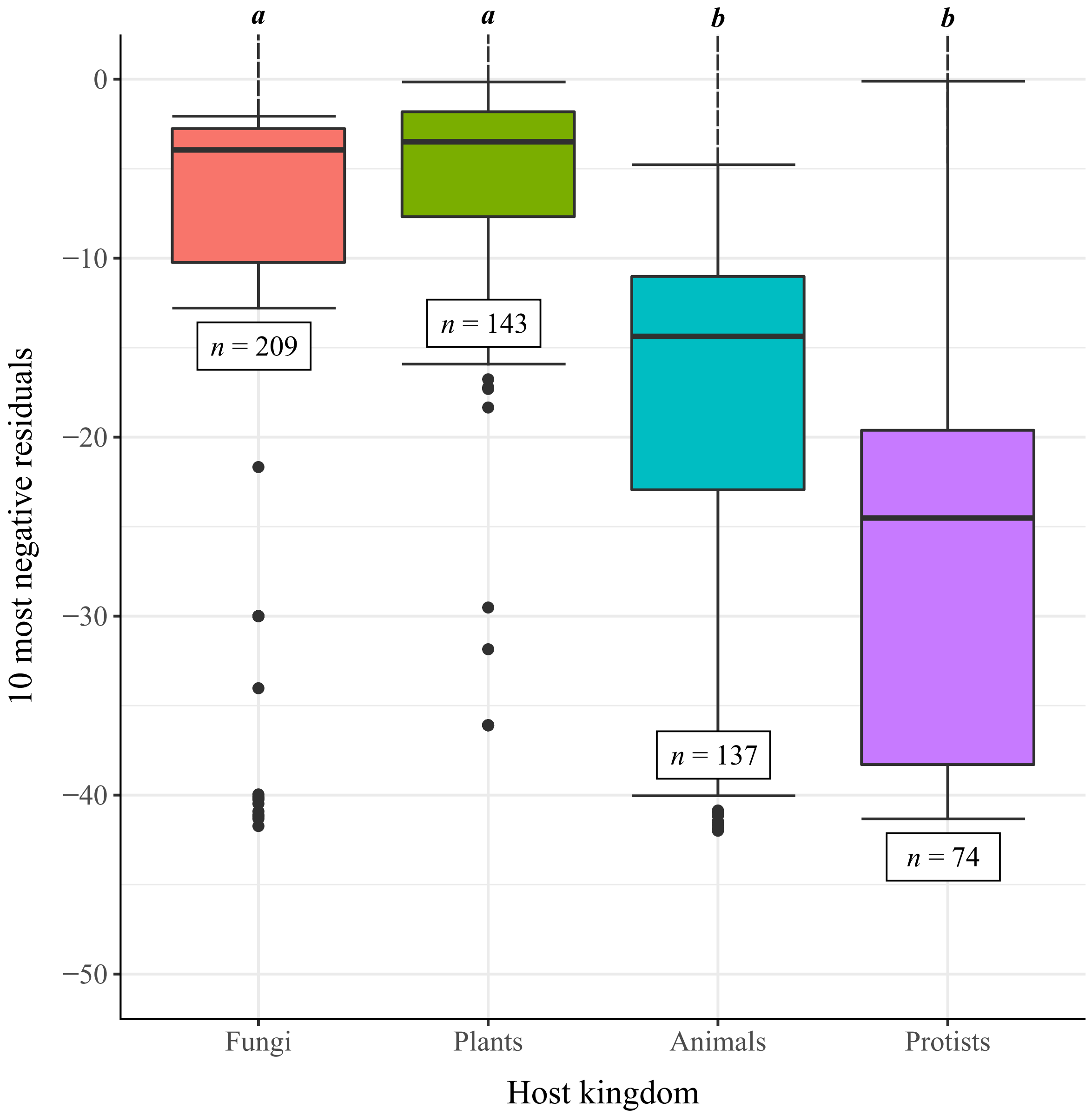


**Figure S11.** Eukaryotic hosts most susceptible to viral spillovers in all samples. The residuals were computed by Random TaPas ($n = 10$ runs) using PACo (Procrustes Approach to Cophylogeny), all samples ($n = 6$) confounded, from which the ten most negative virus/host associations were analysed (total sample size: $n = 10 \times 6 \times 10_{MostNeg} - PosRes = 563$). Significant results (Dunn test, BH correction) are marked with letters $a$ and $b$ ($\alpha = 0.01$). Outliers are shown as dots. Similar results were previously found with the GD model (figure 4).

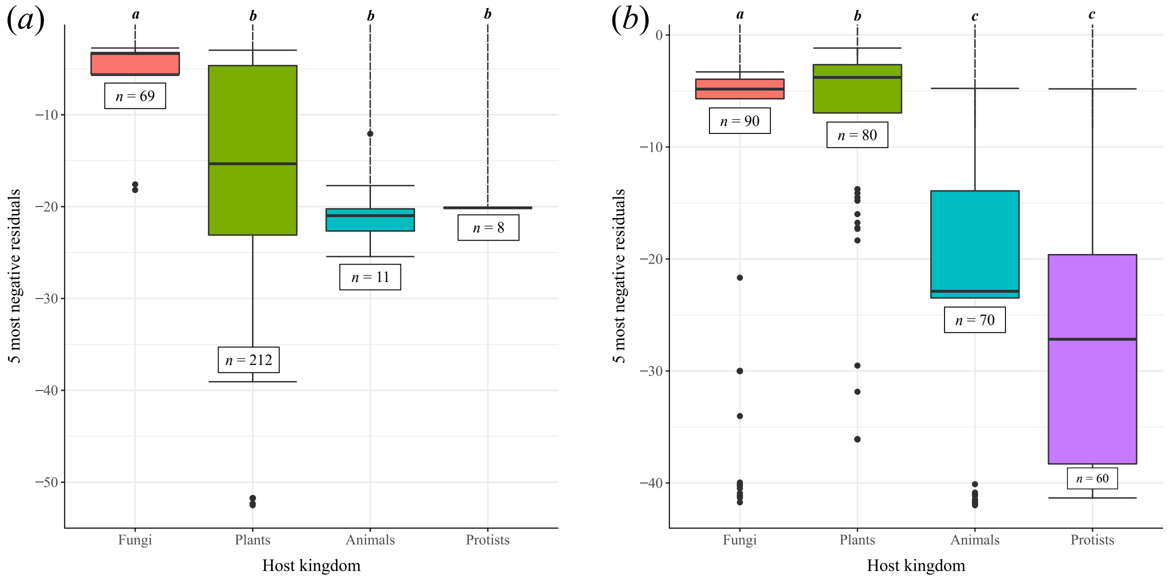


**Figure S12.** Eukaryotic hosts most susceptible to viral spillovers in all samples. The residuals were computed by Random TaPas ($n = 10$ runs) using ($a$) GD and ($b$) PACo, all samples ($n = 6$) confounded, from which the five most negative virus/host associations were analysed (total sample size: $n = 10 \times 6 \times 5_{MostNeg}$ for each model). Significant results (Dunn test, BH correction) are marked with letters from $a$ to $c$ ($\alpha = 0.01$). Outliers are shown as dots.

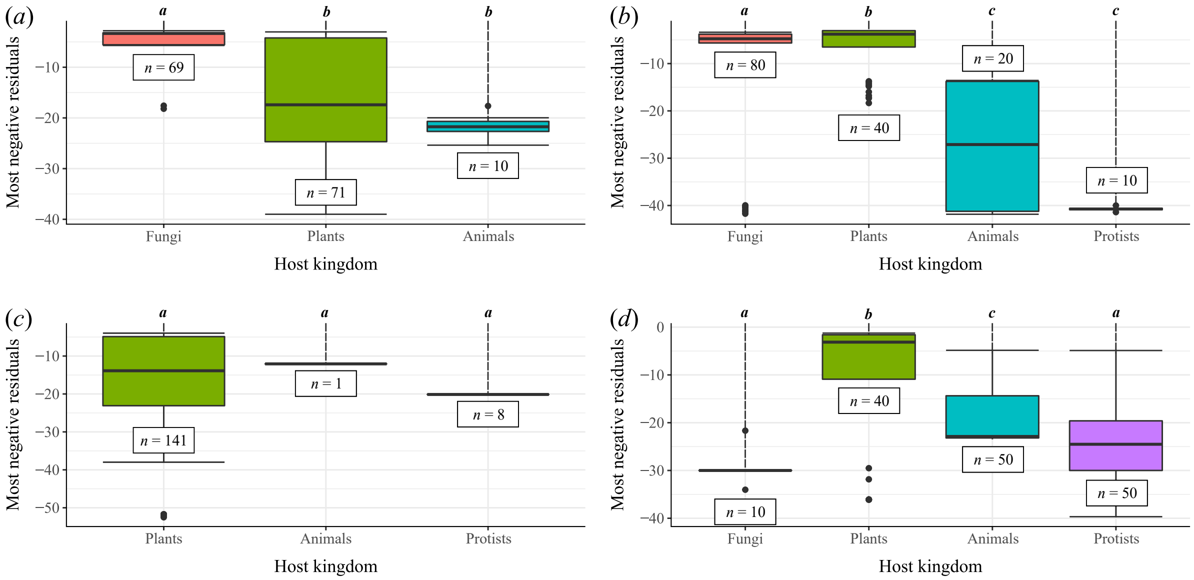


**Figure S13.** Eukaryotic hosts most susceptible to viral spillovers in soil and lake sediments. The residuals were computed by Random TaPas ($n = 10$ runs) using GD and PACo, from which the five most negative virus/host associations were analysed (total sample size: $n = 10 \times 3 \times 5_{MostNeg}$ for each environment/model). ($a$) Soil – GD; ($b$) Soil – PACo; ($c$) Lake sediments – GD; and ($d$) Lake sediments – PACo. Significant results (Dunn test, BH correction) are marked with letters from $a$ to $c$ ($\alpha = 0.01$). Outliers are shown as dots.

**Table S1.** Coordinates and sampling date of Lake Hazen soil and sediment sites. This table was adapted from previous work [22].

| Environment | Site | Coordinates | Sampling date |
|---|---|---|---|
| | C-Soil | 81°79′382″ N; 70°44′486″ W | June 3, 2017 |
| Soil | L-Soil | 81°80′332″ N; 71°54′239″ W | June 7, 2017 |
| | H-Soil | 81°84′840″ N; 70°83′849″ W | June 3, 2017 |
| | C-Sed | 81°80′343″ N; 70°50′447″ W | May 27, 2017 |
| Sediments | L-Sed | 80°80′521″ N; 70°52′699″ W | June 1, 2017 |
| | H-Sed | 81°84′150″ N; 70°85′175″ W | May 24, 2017 |

**Table S2.** RNA bioanalysis for each sample. Quantification of DNA fluorescence assays can be found in previous work [22].

| Environment | Sample | Volume ($\mu$L) | Concentration ($\eta$g/$\mu$L) | Total RNA ($\eta$g) |
|---|---|---|---|---|
| | C-Soil | 48 | 278.34 | 13,360.32 |
| Soil | L-Soil | 48 | 124.76 | 5,988.48 |
| | H-Soil | 48 | 67.59 | 3,244.32 |
| | C-Sed | 48 | 52.56 | 2,522.88 |
| Lake sediments | L-Sed | 48 | 114.52 | 5,496.96 |
| | H-Sed | 48 | 8.52 | 408.96 |

**Table S3.** Taxonomic annotations at the superkingdom level assigned to the DNA contigs assembled with SPAdes and metaSPAdes. They were both tested on a replicate of the L-Soil sample.

| Number of contigs | Assembly pipeline | |
|---|---|---|
| | SPAdes | metaSPAdes |
| Generated | 846,341 | 1,686,159 |
| Classified (%)[a] | 361,821 (42.751%) | 606,751 (35.984%) |
| Classified as "Archaea" (%)[b] | 5,267 (1.456%) | 5,754 (0.948%) |
| Classified as "Bacteria" (%)[b] | 348,591 (96.343%) | 588,910 (97.060%) |
| Classified as "Eukaryota" (%)[b] | 524 (0.145%) | 680 (0.112%) |
| Classified as "Viruses" (%)[b] | 20 (0.006%) | 25 (0.004%) |
| Classified as "Other" (%)[b] | 7,419 (2.050%) | 11,382 (1.876%) |

[a] Proportions were calculated with the number of contigs generated.
[b] Proportions were calculated with the number of classified contigs.

**Table S4.** Taxonomic annotations at the superkingdom level assigned to the RNA contigs assembled with Trinity, rnaSPAdes and metaSPAdes. They were all tested on a replicate of the L-Soil sample.

| Number of contigs | Assembly pipeline | | |
|---|---|---|---|
| | Trinity | rnaSPAdes | metaSPAdes |
| Generated | 727,16 | 1,564,307 | 672,527 |
| Classified (%)[a] | 410,375 (56.44%) | 559,032 (35.74%) | 330,635 (49.16%) |
| Classified as "Archaea" (%)[b] | 2,821 (0.69%) | 3,812 (0.68%) | 2,405 (0.73%) |
| Classified as "Bacteria" (%)[b] | 393,076 (95.79%) | 534,946 (95.69%) | 317,907 (96.15%) |
| Classified as "Eukaryota" (%)[b] | 7,621 (1.86%) | 10,496 (1.88%) | 5,293 (1.60%) |
| Classified as "Viruses" (%)[b] | 177 (0.04%) | 220 (0.04%) | 154 (0.05%) |
| Classified as "Other" (%)[b] | 6,680 (1.63%) | 9,558 (1.71%) | 4,876 (1.47%) |

[a] Proportions were calculated with the number of contigs generated.
[b] Proportions were calculated with the number of classified contigs.

**Table S5.** The average proportions of taxonomic annotations in all samples. The proportions of superkingdoms were calculated for each DNA ($n = 2$ *per* sample) and RNA ($n = 3$ *per* sample) replicate, and averages were then computed per sample based on these values.

| Nucleic acid | Sample | Proportion of classified contigs (%)[a] | Proportions of taxonomic annotations (%)[b] | | | | |
|---|---|---|---|---|---|---|---|
| | | | Archaea | Bacteria | Eukaryota | Viruses | Other |
| DNA | C-Soil | 36.813 | 0.429 | 97.850 | 0.311 | 0.004 | 1.407 |
| | L-Soil | 36.013 | 0.930 | 97.083 | 0.111 | 0.003 | 1.872 |
| | H-Soil | 38.915 | 0.711 | 94.633 | 2.575 | 0.017 | 2.064 |
| | C-Sed | 26.674 | 0.496 | 95.247 | 0.232 | 0.137 | 3.888 |
| | L-Sed | 29.078 | 1.440 | 95.028 | 0.172 | 0.144 | 3.216 |
| | H-Sed | 28.910 | 0.233 | 96.072 | 0.806 | 0.594 | 2.296 |
| RNA | C-Soil | 41.967 | 0.556 | 93.707 | 4.312 | 0.169 | 1.255 |
| | L-Soil | 35.755 | 0.690 | 95.634 | 1.899 | 0.042 | 1.735 |
| | H-Soil | 38.206 | 0.504 | 92.080 | 5.571 | 0.184 | 1.661 |
| | C-Sed | 20.160 | 0.846 | 89.198 | 6.435 | 0.127 | 3.395 |
| | L-Sed | 20.609 | 0.609 | 91.137 | 5.224 | 0.132 | 2.898 |
| | H-Sed | 24.932 | 1.886 | 72.555 | 18.416 | 0.865 | 6.279 |

[a] Proportions were calculated with the number of contigs generated.
[b] Proportions were calculated with the number of classified contigs.

**Table S6.** The number of viral High-scoring Segment Pairs (HSPs) returned by BLASTn and Bowtie2 on the viral contigs, and by BLASTp on the predicted genes. These three alignment tools were tested on the contigs of one RNA replicate of each sample ($n = 6$).

| Replicate of the sample | Number of HSPs returned by alignment tool | | |
|---|---|---|---|
| | BLASTn | Bowtie2 | BLASTp |
| C-Soil | 287 | 0 | 1,179 |
| L-Soil | 178 | 3 | 1,272 |
| H-Soil | 475 | 2 | 1,957 |
| C-Sed | 310 | 0 | 4,582 |
| L-Sed | 829 | 3 | 6,807 |
| H-Sed | 2,468 | 18 | 9,738 |

**Table S7.** Virus/host associations of the mean residuals computed by Random TaPas. The algorithm was run $n = 10$ times *per* sampling site with the global-fit model GD (geodesic distances in tree space), from which mean residuals were calculated.

| | Mean residuals for GD | | | | | |
|---|---|---|---|---|---|---|
| | Control | | Low runoff | | High runoff | |
| | C-Soil | C-Sed | L-Soil | L-Sed | H-Soil | H-Sed |
| **Plant viruses and their hosts** | | | | | | |
| *African oil palm ringspot virus - Elaeis guineensis* | | | 24.74 | | | 8.10 |
| *Alfalfa mosaic virus - Capsicum annuum* | | | 1.19 | | | |
| *Alfalfa mosaic virus - Cicer arietinum* | | | 1.97 | | | |
| *Alfalfa mosaic virus - Lactuca sativa* | | | 3.68 | | | |
| *Alfalfa mosaic virus - Nicotiana tabacum* | | | 0.32 | | | |
| *Alfalfa mosaic virus - Phaseolus vulgaris* | | | 2.90 | | | |
| *Alfalfa mosaic virus - Solanum lycopersicum* | | | -0.78 | | | |
| *Alfalfa mosaic virus - Solanum tuberosum* | | | -0.35 | | | |
| *Apple chlorotic leaf spot virus - Prunus persica* | | | | | | 1.14 |
| *Apple green crinkle associated virus - Malus domestica* | | | 16.58 | -8.10 | | 12.07 |
| *Arhar cryptic virus I - Cajanus cajan* | | -5.27 | | -37.59 | | -51.88 |
| *Asian prunus virus 1 - Prunus persica* | | | | | | 1.99 |
| *Asian prunus virus 2 - Prunus mume* | | | | -19.91 | | 7.49 |
| *Asian prunus virus 2 - Prunus persica* | | | | | | -0.97 |
| *Asian prunus virus 3 - Prunus persica* | | | | | | 1.46 |
| *Bell pepper alphaendornavirus - Capsicum annuum* | 2.37 | -0.10 | 2.31 | | | 5.34 |
| *Cannabis cryptic virus - Cannabis sativa* | | -4.61 | -26.16 | | | -31.91 |
| *Cardamine chlorotic fleck virus - Arabidopsis thaliana* | | | -1.91 | | | |
| *Carnation mottle virus - Malus domestica* | | | | -22.73 | | |
| *Caucasus prunus virus - Prunus dulcis* | | | | | | 10.39 |
| *Cherry green ring mottle virus - Prunus avium* | | | | -5.87 | | -2.21 |
| *Cherry green ring mottle virus - Prunus persica* | | | | | | 0.26 |
| *Cherry mottle leaf virus - Prunus avium* | | | | -3.50 | | 4.30 |
| *Cherry necrotic rusty mottle virus - Prunus avium* | | | | -5.28 | | 4.23 |
| *Cherry necrotic rusty mottle virus - Prunus persica* | | | | | | 3.02 |
| *Cherry rusty mottle associated virus - Prunus avium* | | | | -5.97 | | 1.32 |
| *Cherry twisted leaf associated virus - Prunus avium* | | | | -5.51 | | 0.88 |
| *Cherry virus A - Prunus avium* | | | | -7.15 | | 8.24 |
| *Citrus concave gum associated virus - Citrus sinensis* | | | -16.66 | | -5.04 | |

| | | | | | | |
|---|---|---|---|---|---|---|
| *Cocksfoot mottle virus - Triticum aestivum* | 2.80 | | | | | |
| *Cowpea mild mottle virus - Vigna unguiculata* | | | | | | -9.10 |
| *Cucumis melo alphaendornavirus - Cucumis melo* | | | | | | 6.16 |
| *Galinsoga mosaic virus - Nicotiana tabacum* | | | | | 2.68 | |
| *Grapevine endophyte alphaendornavirus - Vitis vinifera* | | -0.88 | | | | |
| *Grapevine Pinot gris virus - Vitis vinifera* | | | | -12.91 | | |
| *Grapevine rupestris stem pitting-associated virus - Vitis vinifera* | | | | 7.75 | | |
| *Helianthus annuus alphaendornavirus - Helianthus annuus* | | -3.39 | 3.41 | | | -0.96 |
| *Lagenaria siceraria endornavirus Hubei - Benincasa hispida* | | | 28.28 | | | |
| *Lettuce ring necrosis virus - Lactuca sativa* | | | -4.77 | | | |
| *Maize associated totivirus 1 - Zea mays* | | | -5.73 | -12.14 | | -6.32 |
| *Maize associated totivirus 2 - Zea mays* | | | -7.64 | -11.39 | | -11.93 |
| *Maize associated totivirus 3 - Zea mays* | | | -6.62 | -10.55 | | -7.21 |
| *Melon partitivirus - Cucumis melo* | | | -37.45 | | | -14.64 |
| *Melon yellowing-associated virus - Cucumis melo* | | | | | | 18.61 |
| *Mume virus A - Prunus mume* | | | | | | 13.34 |
| *Oryza rufipogon alphaendornavirus - Oryza rufipogon* | | | | | | 10.41 |
| *Oryza sativa alphaendornavirus - Oryza sativa Japonica Group* | 7.03 | -4.88 | 20.13 | | | |
| *Papaya mosaic virus - Carica papaya* | | | 16.91 | | | |
| *Peach chlorotic mottle virus - Prunus persica* | | | | | | 0.06 |
| *Peach mosaic virus - Prunus persica* | | | | | | 1.63 |
| *Pepper cryptic virus 1 - Capsicum annuum* | | -3.28 | 1.47 | | | 0.30 |
| *Pepper cryptic virus 2 - Capsicum annuum* | | -3.35 | 0.69 | | | -2.49 |
| *Pepper virus A - Capsicum annuum* | | | 5.69 | | | -4.71 |
| *Phaseolus vulgaris alphaendornavirus 1 - Phaseolus vulgaris* | | | | | | 15.38 |
| *Phaseolus vulgaris alphaendornavirus 2 - Phaseolus vulgaris* | 8.77 | | 8.56 | | | -7.94 |
| *Potato latent virus - Solanum tuberosum* | | | | | | -2.24 |
| *Potato necrosis virus - Solanum tuberosum* | | | | | 0.88 | 0.58 |
| *Potato virus H - Solanum tuberosum* | | | | | | -6.58 |
| *Potato virus M - Solanum tuberosum* | | | 2.26 | | | -3.41 |
| *Potato virus P - Solanum tuberosum* | | | 6.22 | | | -4.39 |
| *Potato virus S - Solanum tuberosum* | | | | | | 4.07 |
| *Potato virus X - Brassica rapa* | | | 15.99 | | | |
| *Potato virus X - Solanum tuberosum* | | | -0.28 | | | |
| *Raphanus sativas chrysovirus 1 - Raphanus sativus* | | | | | -4.20 | -1.50 |
| *Raphanus sativus cryptic virus 1 - Raphanus sativus* | | | -16.52 | | 13.34 | -7.65 |
| *Raphanus sativus cryptic virus 2 - Raphanus sativus* | | -4.04 | | | | -7.88 |

| | | | | | | |
|---|---|---|---|---|---|---|
| *Raphanus sativus cryptic virus 3 - Raphanus sativus* | | -3.90 | -19.42 | | | -6.05 |
| *Red clover vein mosaic virus - Cicer arietinum* | | | | | | 12.34 |
| *Rottboellia yellow mottle virus - Zea mays* | | | -5.37 | | | |
| *Southern bean mosaic virus - Phaseolus vulgaris* | -2.21 | | 0.08 | | | |
| *Southern cowpea mosaic virus - Glycine max* | 0.68 | | | | | |
| *Southern cowpea mosaic virus - Phaseolus vulgaris* | 0.74 | | 1.08 | | | |
| *Soybean yellow common mosaic virus - Glycine max* | -2.95 | | | | | |
| *Spinach cryptic virus 1 - Spinacia oleracea* | | | -3.18 | | 12.97 | -17.63 |
| *Strawberry mild yellow edge virus - Chenopodium quinoa* | | | | | -2.75 | |
| *Tomato bushy stunt virus - Capsicum annuum* | | | | | 2.03 | |
| *Zea mays chrysovirus 1 - Zea mays* | | | | | | 2.07 |
| **Animal viruses and their hosts** | | | | | | |
| *Akhmeta virus - Homo sapiens* | | | | | | 5.07 |
| *Apis dicistrovirus - Apis mellifera* | | | | -7.67 | | |
| *Banna virus strain JKT-6423 - Homo sapiens* | | | | | | -4.32 |
| *Cotia virus SPAn232 - Mus musculus* | | | | | | 4.96 |
| *Cowpox virus - Callithrix jacchus* | | | 7.65 | | | |
| *Cowpox virus - Canis lupus familiaris* | | | 5.01 | | | |
| *Cowpox virus - Homo sapiens* | | | | | | 3.43 |
| *Cowpox virus - Loxodonta africana* | | | | | | -0.40 |
| *Cowpox virus - Mus musculus* | | | | | | -1.85 |
| *Cowpox virus - Panthera pardus* | | | | | | 2.39 |
| *Cowpox virus - Puma concolor* | | | | | | 1.35 |
| *Cowpox virus - Rattus norvegicus* | | | 6.48 | | | -0.66 |
| *Cowpox virus - Sorex araneus* | | | | | | 0.91 |
| *Ectromelia virus - Mus musculus* | | | | | | 7.05 |
| *Equid gammaherpesvirus 5 - Equus caballus* | | | | 44.13 | | |
| *Human alphaherpesvirus 3 - Homo sapiens* | | | | 1.87 | | |
| *Human gammaherpesvirus 8 - Homo sapiens* | | | | 39.08 | | |
| *Laurel Lake virus - Ixodes scapularis* | | | -21.69 | | | |
| *Liao ning virus - Mus musculus* | | | | | | -8.16 |
| *Monkeypox virus Zaire 96-I-16 - Homo sapiens* | | | | | | 3.89 |
| *Monkeypox virus Zaire 96-I-16 - Mus musculus* | | | | | | 1.98 |
| *Nodamura virus - Aedes aegypti* | | | | 0.81 | | |
| *Nodamura virus - Aedes albopictus* | | | | | | -8.29 |
| *Nodamura virus - Apis mellifera* | | | | 11.58 | | |
| *NY_014 poxvirus - Homo sapiens* | | | | | | -1.85 |

| | | | | | | |
|---|---|---|---|---|---|---|
| *Penaeid shrimp infectious myonecrosis virus - Penaeus vannamei* | | 15.97 | | | | -12.76 |
| *Shrimp hemocyte iridescent virus - Penaeus vannamei* | | | | | | 7.21 |
| *Vaccinia virus - Homo sapiens* | | | | | | 2.58 |
| *Yaba-like disease virus - Homo sapiens* | | | | | | 3.10 |
| *Yaba monkey tumor virus - Homo sapiens* | | | | | | 0.30 |
| *Yokapox virus - Mus musculus* | | | | | | 0.24 |
| *Yongsan tombus-like virus 1 - Aedes albopictus* | | | | | | -5.02 |
| **Fungal viruses and their hosts** | | | | | | |
| *Colletotrichum gloeosporioides chrysovirus 1 - Colletotrichum gloeosporioides* | | | -13.83 | | -5.60 | |
| *Erysiphe necator mitovirus 1 - Erysiphe necator* | | | | | | 4.81 |
| *Erysiphe necator mitovirus 2 - Erysiphe necator* | | | | | | 10.63 |
| *Erysiphe necator mitovirus 3 - Erysiphe necator* | | | | | 0.14 | |
| *Fusarium poae mitovirus 3 - Fusarium poae* | | | | | 0.53 | |
| *Fusarium poae mitovirus 4 - Fusarium poae* | | | | | 0.79 | |
| *Fusarium poae narnavirus 2 - Fusarium poae* | | | | | -2.72 | |
| *Fusarium poae negative-stranded virus 1 - Fusarium poae* | | | | | -2.70 | |
| *Fusarium poae victorivirus 1 - Fusarium poae* | | | | | 2.55 | |
| *Fusarium poae virus 1 - Fusarium poae* | -3.31 | | | | | |
| *Fusarium poae virus 1-240374 - Fusarium poae* | -3.27 | | | | | |
| *Pleurotus ostreatus virus 1 - Pleurotus ostreatus* | -3.95 | | | 48.76 | | |
| *Pseudogymnoascus destructans partitivirus-pa - Pseudogymnoascus destructans* | | | | | -5.66 | |
| *Rhizoctonia mitovirus 1 - Rhizoctonia solani* | 9.07 | | | | | |
| *Rhizoctonia solani dsRNA virus 3 - Rhizoctonia solani* | | | | | | 2.78 |
| *Rhizoctonia solani dsRNA virus 4 - Rhizoctonia solani* | | | -1.84 | | | 0.27 |
| *Rhizoctonia solani endornavirus 1 - Rhizoctonia solani* | -2.78 | | -10.12 | | | -1.72 |
| *Rhizoctonia solani endornavirus 2 - Rhizoctonia solani* | -2.78 | | -9.96 | | | -8.40 |
| *Rhizoctonia solani virus 717 - Rhizoctonia solani* | | | -1.74 | | | |
| *Ustilaginoidea virens RNA virus 3 - Ustilaginoidea virens* | | | | | 2.76 | |
| **Protist viruses and their hosts** | | | | | | |
| *Acanthocystis turfacea chlorella virus 1 - Chlorella heliozoae* | | | | | | 5.55 |
| *Asterionellopsis glacialis RNA virus - Asterionellopsis glacialis* | | | | 34.62 | | 2.29 |
| *Aureococcus anophagefferens virus - Aureococcus anophagefferens* | | | | | | 4.33 |
| *Cedratvirus A11 - Acanthamoeba castellanii str. Neff* | | | 28.42 | | | |
| *Heterosigma akashiwo virus 01 - Heterosigma akashiwo* | | | | | | 3.45 |
| *Only Syngen Nebraska virus 5 - Chlorella variabilis* | | 1.98 | | -6.28 | | |
| *Ostreococcus tauri virus 1 - Ostreococcus tauri* | | | | | | 7.82 |
| *Ostreococcus tauri virus 2 - Ostreococcus tauri* | | | | | | 4.76 |

| | | | |
|---|---|---|---|
| *Ostreococcus tauri virus OtV5 - Ostreococcus tauri* | | | 5.93 |
| *Pacmanvirus A23 - Acanthamoeba castellanii str. Neff* | 18.64 | | 20.13 |
| *Paramecium bursaria Chlorella virus 1 - Chlorella variabilis* | | 3.53 | -4.14 |
| *Paramecium bursaria Chlorella virus AR158 - Chlorella variabilis* | 1.85 | -5.33 | 6.15 |
| *Paramecium bursaria Chlorella virus CVA-1 - Micractinium conductrix* | | | 3.39 |
| *Paramecium bursaria Chlorella virus FR483 - Micractinium conductrix* | | | 2.76 |
| *Paramecium bursaria Chlorella virus NY2A - Chlorella variabilis* | 2.01 | -3.00 | |
| *Paramecium bursaria Chlorella virus NYs1 - Chlorella variabilis* | 0.76 | 2.07 | |
| *Phaeocystis globosa virus - Phaeocystis globosa* | | | -17.53 |

**Table S8.** Virus-host associations of the mean residuals computed by Random TaPas. The algorithm was run $n = 10$ times *per* sampling site with the global-fit model PACo (Procrustes Approach to Cophylogeny), from which mean residuals were calculated.

| | Mean residuals for PACo | | | | | |
|---|---|---|---|---|---|---|
| | Control | | Low runoff | | High runoff | |
| | C-Soil | C-Sed | L-Soil | L-Sed | H-Soil | H-Sed |
| **Plant viruses and their hosts** | | | | | | |
| *African oil palm ringspot virus - Elaeis guineensis* | | | -2.11 | | | 14.12 |
| *Alfalfa mosaic virus - Capsicum annuum* | | | 1.25 | | | |
| *Alfalfa mosaic virus - Cicer arietinum* | | | 3.44 | | | |
| *Alfalfa mosaic virus - Lactuca sativa* | | | 1.66 | | | |
| *Alfalfa mosaic virus - Nicotiana tabacum* | | | 2.98 | | | |
| *Alfalfa mosaic virus - Phaseolus vulgaris* | | | 1.06 | | | |
| *Alfalfa mosaic virus - Solanum lycopersicum* | | | 3.61 | | | |
| *Alfalfa mosaic virus - Solanum tuberosum* | | | 1.06 | | | |
| *Apple chlorotic leaf spot virus - Prunus persica* | | | | | | 0.49 |
| *Apple green crinkle associated virus - Malus domestica* | | | 12.94 | 34.85 | | 22.22 |
| *Arhar cryptic virus I - Cajanus cajan* | | 5.81 | | -6.90 | | -35.01 |
| *Asian prunus virus 1 - Prunus persica* | | | | | | 5.10 |
| *Asian prunus virus 2 - Prunus mume* | | | | 33.53 | | 8.68 |
| *Asian prunus virus 2 - Prunus persica* | | | | | | 1.83 |
| *Asian prunus virus 3 - Prunus persica* | | | | | | 4.53 |
| *Bell pepper alphaendornavirus - Capsicum annuum* | 12.77 | 2.11 | 1.36 | | | 9.44 |
| *Cannabis cryptic virus - Cannabis sativa* | | -4.47 | 21.59 | | | -0.03 |
| *Cardamine chlorotic fleck virus - Arabidopsis thaliana* | | | 19.58 | | | |
| *Carnation mottle virus - Malus domestica* | | | | -5.07 | | |
| *Caucasus prunus virus - Prunus dulcis* | | | | | | 19.10 |
| *Cherry green ring mottle virus - Prunus avium* | | | | 7.71 | | 0.53 |
| *Cherry green ring mottle virus - Prunus persica* | | | | | | 3.84 |
| *Cherry mottle leaf virus - Prunus avium* | | | | -2.47 | | 0.03 |
| *Cherry necrotic rusty mottle virus - Prunus avium* | | | | 2.23 | | 4.00 |
| *Cherry necrotic rusty mottle virus - Prunus persica* | | | | | | 3.48 |
| *Cherry rusty mottle associated virus - Prunus avium* | | | | 9.28 | | 0.03 |
| *Cherry twisted leaf associated virus - Prunus avium* | | | | 5.77 | | 6.98 |
| *Cherry virus A - Prunus avium* | | | | -0.43 | | 12.42 |
| *Citrus concave gum associated virus - Citrus sinensis* | | | 31.68 | | 3.86 | |

| | | | | | | |
|---|---|---|---|---|---|---|
| *Cocksfoot mottle virus - Triticum aestivum* | -4.08 | | | | | |
| *Cowpea mild mottle virus - Vigna unguiculata* | | | | | | 5.76 |
| *Cucumis melo alphaendornavirus - Cucumis melo* | | | | | | 11.93 |
| *Galinsoga mosaic virus - Nicotiana tabacum* | | | | | 14.49 | |
| *Grapevine endophyte alphaendornavirus - Vitis vinifera* | | 9.12 | | | | |
| *Grapevine Pinot gris virus - Vitis vinifera* | | | | 10.35 | | |
| *Grapevine rupestris stem pitting-associated virus - Vitis vinifera* | | | | 33.21 | | |
| *Helianthus annuus alphaendornavirus - Helianthus annuus* | | 8.20 | 11.51 | | | 6.74 |
| *Lagenaria siceraria endornavirus Hubei - Benincasa hispida* | | | 13.47 | | | |
| *Lettuce ring necrosis virus - Lactuca sativa* | | | 17.81 | | | |
| *Maize associated totivirus 1 - Zea mays* | | | 1.63 | 0.46 | | 4.88 |
| *Maize associated totivirus 2 - Zea mays* | | | 5.26 | -2.65 | | -1.74 |
| *Maize associated totivirus 3 - Zea mays* | | | 1.93 | 0.32 | | 1.26 |
| *Melon partitivirus - Cucumis melo* | | | -13.26 | | | -5.87 |
| *Melon yellowing-associated virus - Cucumis melo* | | | | | | 14.72 |
| *Mume virus A - Prunus mume* | | | | | | 15.41 |
| *Oryza rufipogon alphaendornavirus - Oryza rufipogon* | | | | | | 12.92 |
| *Oryza sativa alphaendornavirus - Oryza sativa Japonica Group* | -2.94 | -1.29 | -8.06 | | | |
| *Papaya mosaic virus - Carica papaya* | | | 27.13 | | | |
| *Peach chlorotic mottle virus - Prunus persica* | | | | | | 2.78 |
| *Peach mosaic virus - Prunus persica* | | | | | | 3.77 |
| *Pepper cryptic virus 1 - Capsicum annuum* | | -0.50 | 1.56 | | | 2.57 |
| *Pepper cryptic virus 2 - Capsicum annuum* | | 0.99 | 0.61 | | | 2.05 |
| *Pepper virus A - Capsicum annuum* | | | 10.68 | | | -3.32 |
| *Phaseolus vulgaris alphaendornavirus 1 - Phaseolus vulgaris* | | | | | | 14.60 |
| *Phaseolus vulgaris alphaendornavirus 2 - Phaseolus vulgaris* | -1.92 | | -0.06 | | | -4.97 |
| *Potato latent virus - Solanum tuberosum* | | | | | | 4.64 |
| *Potato necrosis virus - Solanum tuberosum* | | | | | 11.24 | 13.71 |
| *Potato virus H - Solanum tuberosum* | | | | | | -5.05 |
| *Potato virus M - Solanum tuberosum* | | | 7.01 | | | -4.98 |
| *Potato virus P - Solanum tuberosum* | | | 6.88 | | | -3.06 |
| *Potato virus S - Solanum tuberosum* | | | | | | 3.18 |
| *Potato virus X - Brassica rapa* | | | 9.04 | | | |
| *Potato virus X - Solanum tuberosum* | | | 0.83 | | | |
| *Raphanus sativas chrysovirus 1 - Raphanus sativus* | | | | | 6.84 | 0.92 |
| *Raphanus sativus cryptic virus 1 - Raphanus sativus* | | | -7.80 | | -1.02 | 0.32 |
| *Raphanus sativus cryptic virus 2 - Raphanus sativus* | | -1.79 | | | | 2.78 |

| | | | | | | |
|---|---|---|---|---|---|---|
| *Raphanus sativus cryptic virus 3 - Raphanus sativus* | | -0.67 | -13.08 | | | 2.16 |
| *Red clover vein mosaic virus - Cicer arietinum* | | | | | | 20.00 |
| *Rottboellia yellow mottle virus - Zea mays* | | | 0.49 | | | |
| *Southern bean mosaic virus - Phaseolus vulgaris* | 9.42 | | 10.09 | | | |
| *Southern cowpea mosaic virus - Glycine max* | 3.39 | | | | | |
| *Southern cowpea mosaic virus - Phaseolus vulgaris* | 2.60 | | 11.28 | | | |
| *Soybean yellow common mosaic virus - Glycine max* | 7.48 | | | | | |
| *Spinach cryptic virus 1 - Spinacia oleracea* | | | 8.95 | | 0.76 | 12.43 |
| *Strawberry mild yellow edge virus - Chenopodium quinoa* | | | | | -3.26 | |
| *Tomato bushy stunt virus - Capsicum annuum* | | | | | 9.96 | |
| *Zea mays chrysovirus 1 - Zea mays* | | | | | | 9.27 |
| **Animal viruses and their hosts** | | | | | | |
| *Akhmeta virus - Homo sapiens* | | | | | | 3.68 |
| *Apis dicistrovirus - Apis mellifera* | | | | -22.91 | | |
| *Banna virus strain JKT-6423 - Homo sapiens* | | | | | | -5.16 |
| *Cotia virus SPAn232 - Mus musculus* | | | | | | -0.31 |
| *Cowpox virus - Callithrix jacchus* | | | -13.55 | | | |
| *Cowpox virus - Canis lupus familiaris* | | | -13.58 | | | |
| *Cowpox virus - Homo sapiens* | | | | | | -0.39 |
| *Cowpox virus - Loxodonta africana* | | | | | | -1.22 |
| *Cowpox virus - Mus musculus* | | | | | | 0.02 |
| *Cowpox virus - Panthera pardus* | | | | | | 3.74 |
| *Cowpox virus - Puma concolor* | | | | | | 1.30 |
| *Cowpox virus - Rattus norvegicus* | | | -13.42 | | | 3.39 |
| *Cowpox virus - Sorex araneus* | | | | | | -1.56 |
| *Ectromelia virus - Mus musculus* | | | | | | 5.14 |
| *Equid gammaherpesvirus 5 - Equus caballus* | | | | -10.23 | | |
| *Human alphaherpesvirus 3 - Homo sapiens* | | | | -5.78 | | |
| *Human gammaherpesvirus 8 - Homo sapiens* | | | | -4.36 | | |
| *Laurel Lake virus - Ixodes scapularis* | | | -41.28 | | | |
| *Liao ning virus - Mus musculus* | | | | | | -8.57 |
| *Monkeypox virus Zaire 96-I-16 - Homo sapiens* | | | | | | 0.92 |
| *Monkeypox virus Zaire 96-I-16 - Mus musculus* | | | | | | 1.22 |
| *Nodamura virus - Aedes aegypti* | | | | -23.25 | | |
| *Nodamura virus - Aedes albopictus* | | | | | | -23.07 |
| *Nodamura virus - Apis mellifera* | | | | -14.68 | | |
| *NY_014 poxvirus - Homo sapiens* | | | | | | -1.24 |

| | | | | | | |
|---|---|---|---|---|---|---|
| *Penaeid shrimp infectious myonecrosis virus - Penaeus vannamei* | | -4.75 | | | | -21.40 |
| *Shrimp hemocyte iridescent virus - Penaeus vannamei* | | | | | | 2.64 |
| *Vaccinia virus - Homo sapiens* | | | | | | 0.86 |
| *Yaba-like disease virus - Homo sapiens* | | | | | | 0.61 |
| *Yaba monkey tumor virus - Homo sapiens* | | | | | | -0.44 |
| *Yokapox virus - Mus musculus* | | | | | | 1.19 |
| *Yongsan tombus-like virus 1 - Aedes albopictus* | | | | | | -22.75 |
| **Fungal viruses and their hosts** | | | | | | |
| *Colletotrichum gloeosporioides chrysovirus 1 - Colletotrichum gloeosporioides* | | | -40.72 | | -4.74 | |
| *Erysiphe necator mitovirus 1 - Erysiphe necator* | | | | | | -10.15 |
| *Erysiphe necator mitovirus 2 - Erysiphe necator* | | | | | | -9.49 |
| *Erysiphe necator mitovirus 3 - Erysiphe necator* | | | | | -5.65 | |
| *Fusarium poae mitovirus 3 - Fusarium poae* | | | | | -2.73 | |
| *Fusarium poae mitovirus 4 - Fusarium poae* | | | | | -2.69 | |
| *Fusarium poae narnavirus 2 - Fusarium poae* | | | | | -2.25 | |
| *Fusarium poae negative-stranded virus 1 - Fusarium poae* | | | | | -2.33 | |
| *Fusarium poae victorivirus 1 - Fusarium poae* | | | | | -2.73 | |
| *Fusarium poae virus 1 - Fusarium poae* | -3.32 | | | | | |
| *Fusarium poae virus 1-240374 - Fusarium poae* | -3.27 | | | | | |
| *Pleurotus ostreatus virus 1 - Pleurotus ostreatus* | -3.96 | | | -29.57 | | |
| *Pseudogymnoascus destructans partitivirus-pa - Pseudogymnoascus destructans* | | | | | -4.80 | |
| *Rhizoctonia mitovirus 1 - Rhizoctonia solani* | -2.02 | | | | | |
| *Rhizoctonia solani dsRNA virus 3 - Rhizoctonia solani* | | | | | | -7.41 |
| *Rhizoctonia solani dsRNA virus 4 - Rhizoctonia solani* | | | -10.11 | | | -10.40 |
| *Rhizoctonia solani endornavirus 1 - Rhizoctonia solani* | -2.78 | | -10.26 | | | -12.55 |
| *Rhizoctonia solani endornavirus 2 - Rhizoctonia solani* | -2.78 | | -10.12 | | | -12.70 |
| *Rhizoctonia solani virus 717 - Rhizoctonia solani* | | | -10.20 | | | |
| *Ustilaginoidea virens RNA virus 3 - Ustilaginoidea virens* | | | | | -5.61 | |
| **Protist viruses and their hosts** | | | | | | |
| *Acanthocystis turfacea chlorella virus 1 - Chlorella heliozoae* | | | | | | -2.78 |
| *Asterionellopsis glacialis RNA virus - Asterionellopsis glacialis* | | | | -19.29 | | -38.51 |
| *Aureococcus anophagefferens virus - Aureococcus anophagefferens* | | | | | | -27.65 |
| *Cedratvirus A11 - Acanthamoeba castellanii str. Neff* | | | -40.75 | | | |
| *Heterosigma akashiwo virus 01 - Heterosigma akashiwo* | | | | | | -26.18 |
| *Only Syngen Nebraska virus 5 - Chlorella variabilis* | | 0.58 | | 5.16 | | |
| *Ostreococcus tauri virus 1 - Ostreococcus tauri* | | | | | | 6.46 |
| *Ostreococcus tauri virus 2 - Ostreococcus tauri* | | | | | | 10.97 |

| | | | |
|---|---|---|---|
| *Ostreococcus tauri virus OtV5 - Ostreococcus tauri* | | | -1.14 |
| *Pacmanvirus A23 - Acanthamoeba castellanii str. Neff* | -4.77 | | 5.11 |
| *Paramecium bursaria Chlorella virus 1 - Chlorella variabilis* | | 1.37 | -2.28 |
| *Paramecium bursaria Chlorella virus AR158 - Chlorella variabilis* | 0.49 | 1.85 | -1.97 |
| *Paramecium bursaria Chlorella virus CVA-1 - Micractinium conductrix* | | | -0.92 |
| *Paramecium bursaria Chlorella virus FR483 - Micractinium conductrix* | | | 2.07 |
| *Paramecium bursaria Chlorella virus NY2A - Chlorella variabilis* | 0.15 | 4.41 | |
| *Paramecium bursaria Chlorella virus NYs1 - Chlorella variabilis* | 0.07 | -1.55 | |
| *Phaeocystis globosa virus - Phaeocystis globosa* | | | -20.64 |
